## Supplementary Table S2 and Figure S1-S25 for "Computational screening of the effects of mutations on protein-protein off-rates and dissociation mechanisms by τRAMD"

**Contents:**

Table S1 – see accompanying Excel file.

Table S2,

Figures S1-S25.

**Table S1: Experimental and computed values of kinetic parameters.** These are provided in the accompanying Excel file**.**

**Table S2. List of possible PP-REs.** For every atom type, the corresponding amino acid atoms are listed together with the type of interaction they are involved in, and the corresponding distance measured.

| Atom type | Protein atoms | Type of interaction | Measure |
|---|---|---|---|
| IP (Positive ionizable) | NH* NZ in ARG, LYS, or HD HE in HIP | Cationic/anionic (IP-IN and IN-IP) | |
| IN (Negative ionizable) | OE* OD* in ASP, GLU | | |
| HD (donor) | O* N* in TYR, SER, LYS, GLN, ARG, HIS, HIE, HID, ASN, THR, and not backbone | H-Bond donor/acceptor (HD-HA and HA-HD) | |
| HA (acceptor) | O* in GLU, ASP, GLN, and not backbone | | |
| AR (aromatic) | CZ* CD* CE* CG* CH* NE* ND* in PHE TRP TYR HIS HIP HIE HID | Aromatic (AR-AR) | Distance between the center of mass (COM) of the atoms involved in the specified interaction |
| HY (hydrophobic) | C* S* in protein, and not CG in ASN ASP, and not CD in GLU GLN ARG, and not CZ in TYR ARG, and not CE in LYS, and not CB in SER THR, and not backbone | Hydrophobic (HY-HY) | |
| BB | backbone | H-Bond donor/acceptor (BB-BB, BB_P-HA, BB_N-HD and HA-BB_P, HD-BB_N) | |
| BB_P | N in backbone | | |
| BB_N | O in backbone | | |
| C | C* and not backbone[1] | Hydrophobic | |

[1]For glycine only, the backbone atoms are considered for this interaction

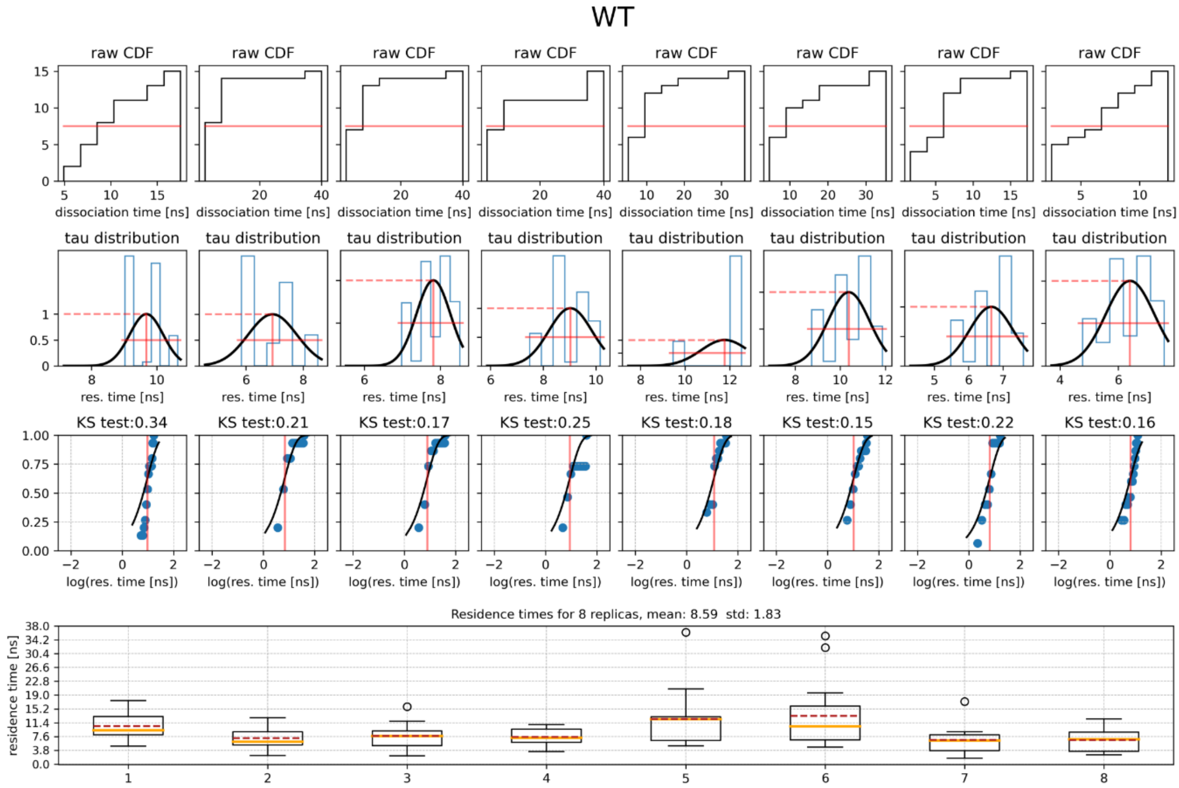
WT
raw CDF
dissociation time [ns]
tau distribution
res. time [ns]
KS test:0.34
KS test:0.21
KS test:0.17
KS test:0.25
KS test:0.18
KS test:0.15
KS test:0.22
KS test:0.16
log(res. time [ns])
Residence times for 8 replicas, mean: 8.59  std: 1.83
residence time [ns]

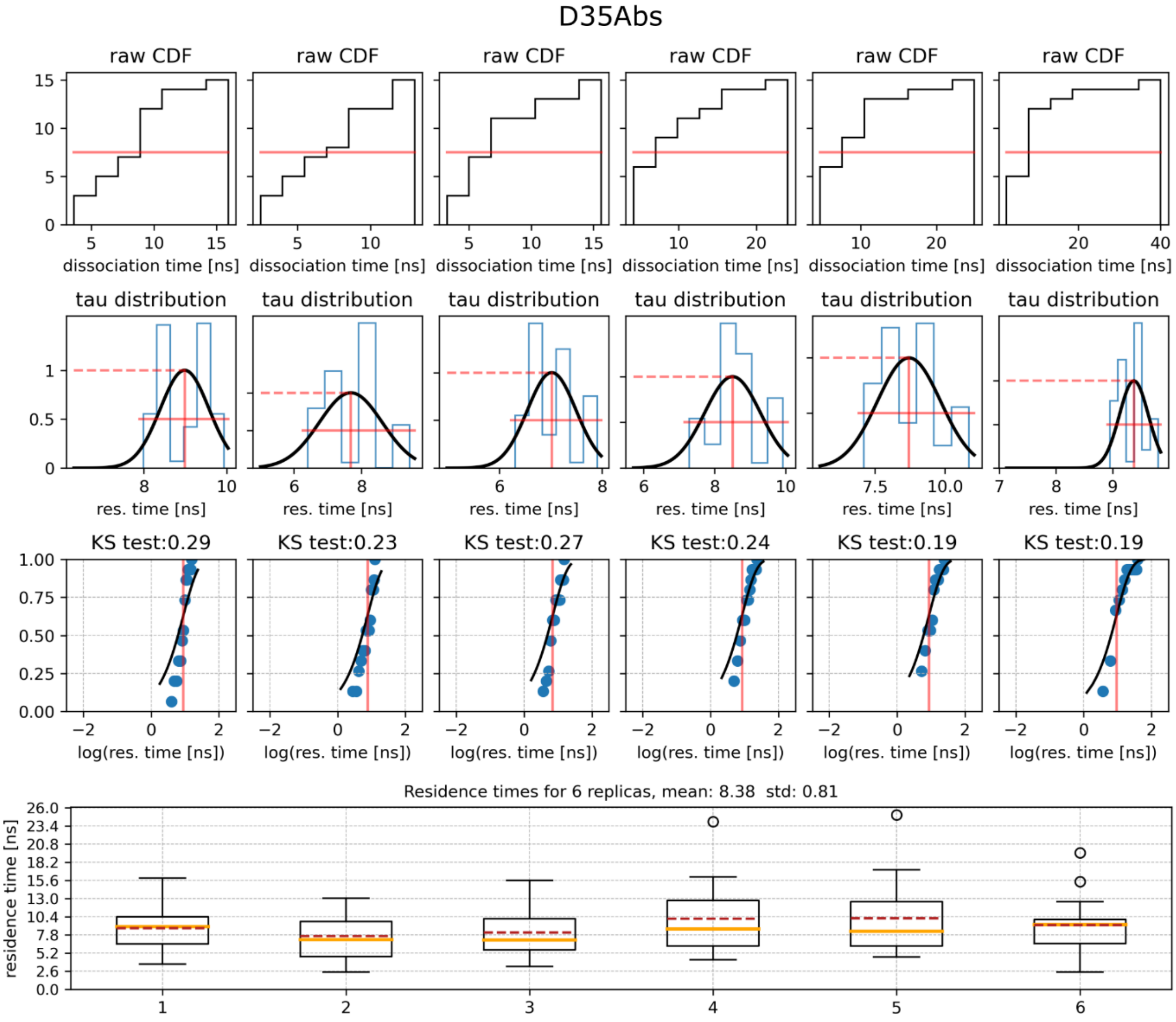
D35Abs
raw CDF
dissociation time [ns]
tau distribution
res. time [ns]
KS test:0.29
KS test:0.23
KS test:0.27
KS test:0.24
KS test:0.19
KS test:0.19
log(res. time [ns])
Residence times for 6 replicas, mean: 8.38  std: 0.81
residence time [ns]

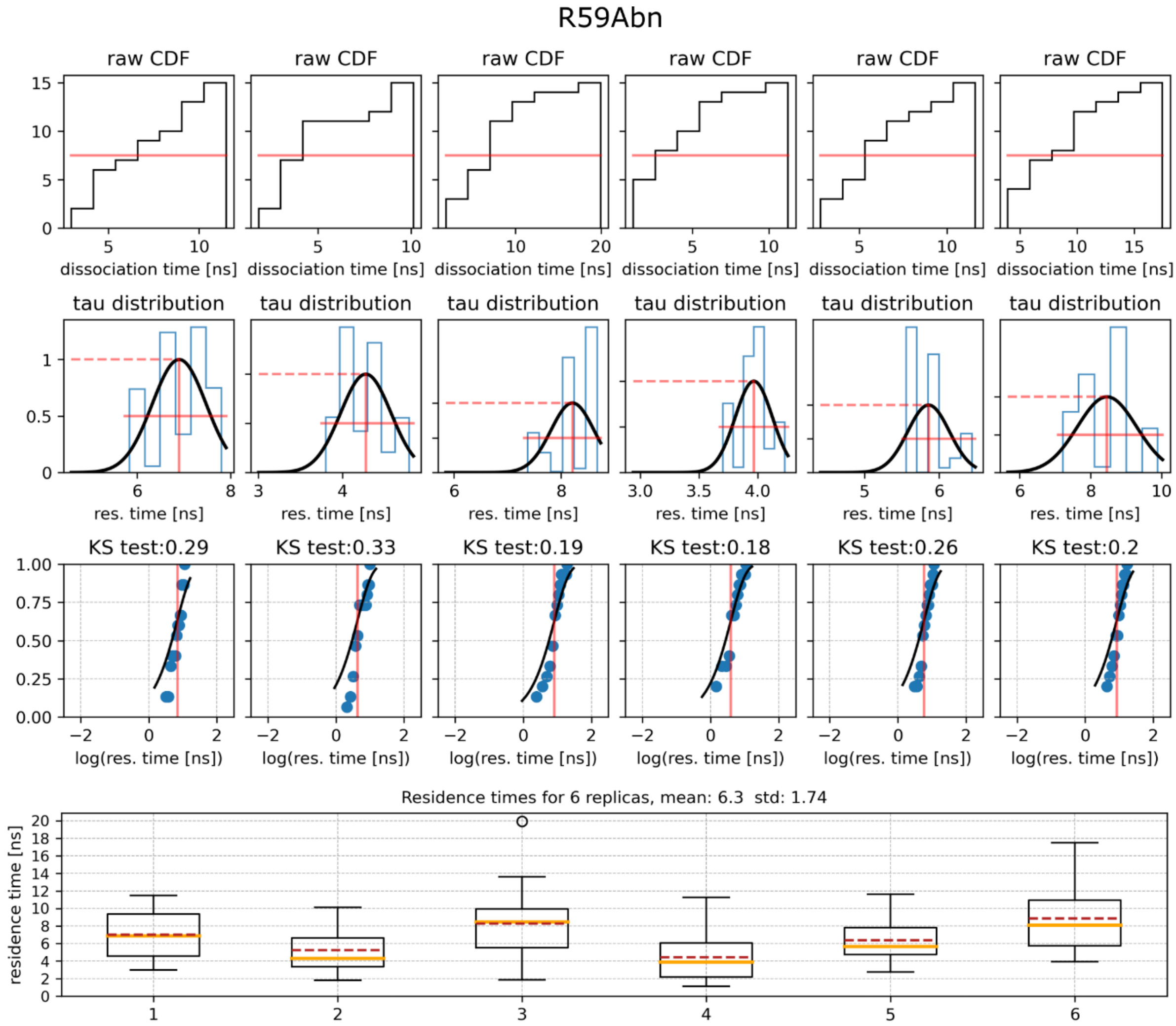


**Figure S1. Statistics of RAMD dissociation trajectories for the WT Bn-Bs complex and two representative mutants.** The plots are shown for WT, D35Abs and R59Abn Bn-Bs complexes using the COM-COM method for deriving the residence time with a random force magnitude of 19 kcal/mol Å. For each mutant, the following four plots are shown: (1) Cumulative distribution function (CDF) for each set of RAMD trajectories (i.e., set of trajectories originating from the same replica). The effective residence time corresponds to 50% of the CDF and is shown by the red solid line. (2) Distribution of the effective residence time for each replica ($\tau_{repl}$) resulting from bootstrapping of the raw data. The corresponding Gaussian distribution is outlined by the black line and the mean and the half-width of each distribution by red lines. (3) Kolmogorov-Smirnov (KS) test to measure the distance between the Poisson cumulative distribution function (PCDF, black line) and the empirical cumulative density function (ECDF, blue points). (4) Boxplot of each $\tau_{repl}$ where median and mean are shown by orange and red dashed lines, respectively, the box extends from the lower to upper quartile values of the data, the whiskers show the range of the data, and the outliers are shown as black circles. The average residence time ($\tau_{RAMD}$) and $SD_{RAMD}$ are indicated above the boxplot.

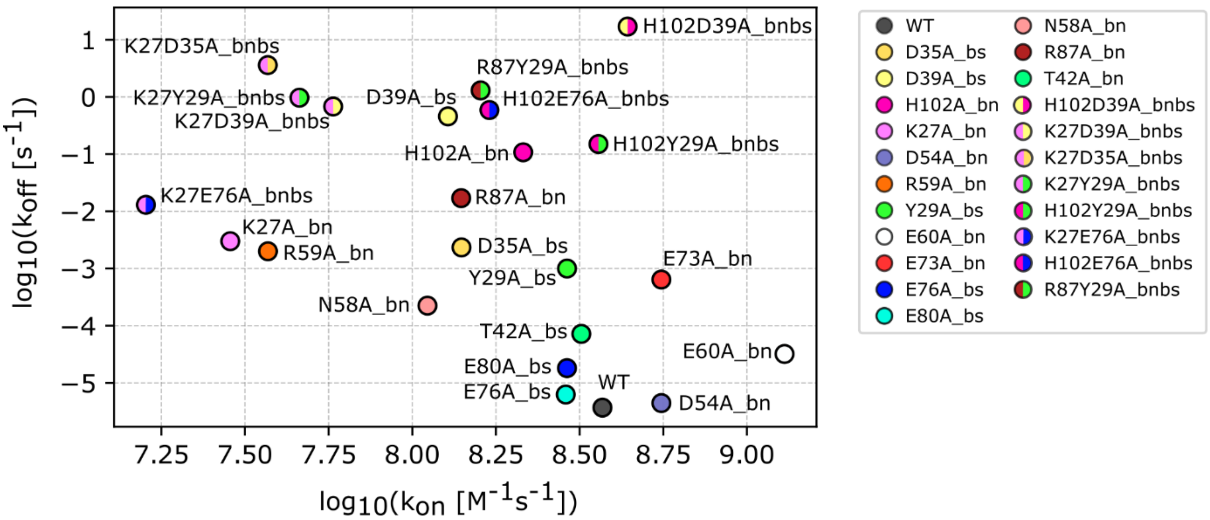


**Figure S2. Normalized experimental kinetic parameters for Bn-BS WT and mutants.** $k_{on}$ and $k_{off}$ values after normalization of the three differently derived sets of experimental values (see Methods for details). A different color is assigned to each single mutant while double mutants are denoted with two colors, each having the same color as the corresponding single mutant.

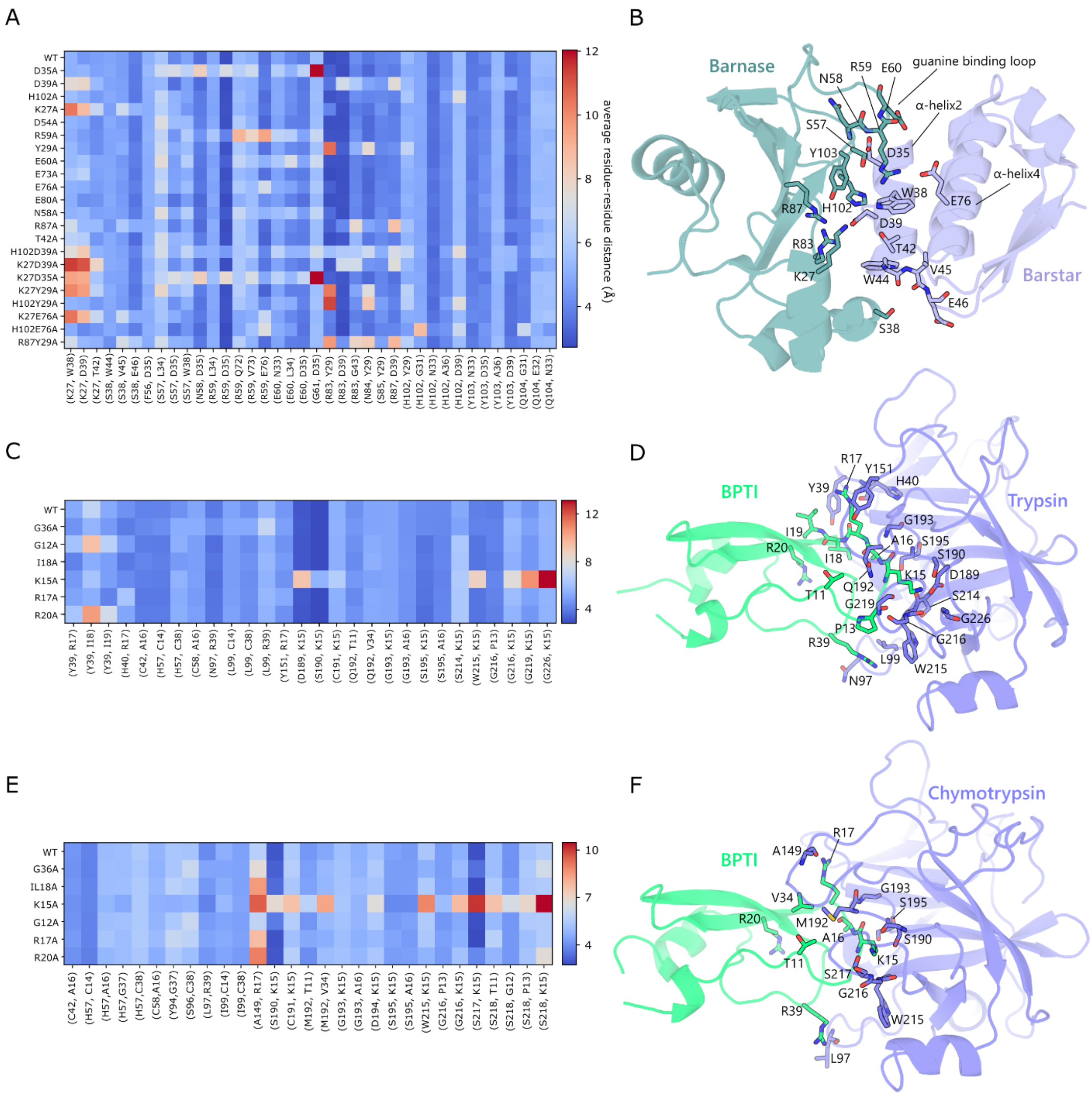


**Figure S3. Interprotein contacts during the equilibration simulations.** The binding interfaces of the complexes are generally well conserved amongst wild-type and mutant complexes with the mutations mostly resulting in the loss of only a few contacts that are directly affected by the mutations themselves. Heatmaps and structures are shown for the **(A, B)** barstar-barnase, **(C, D)** β-trypsin-trypsin inhibitor, and **(E,F)** α-chymotrypsin-trypsin inhibitor complexes The heatmaps display the residue-residue binding site contacts (columns) for the wild-type and all the mutants (rows) averaged over all the equilibration trajectories. Heatmaps are colored on a blue-red scale indicating the average distance between the pair of residues in Å. The corresponding WT complex of each system is shown in cartoon representation, with the main interacting residues shown in sticks and labeled.

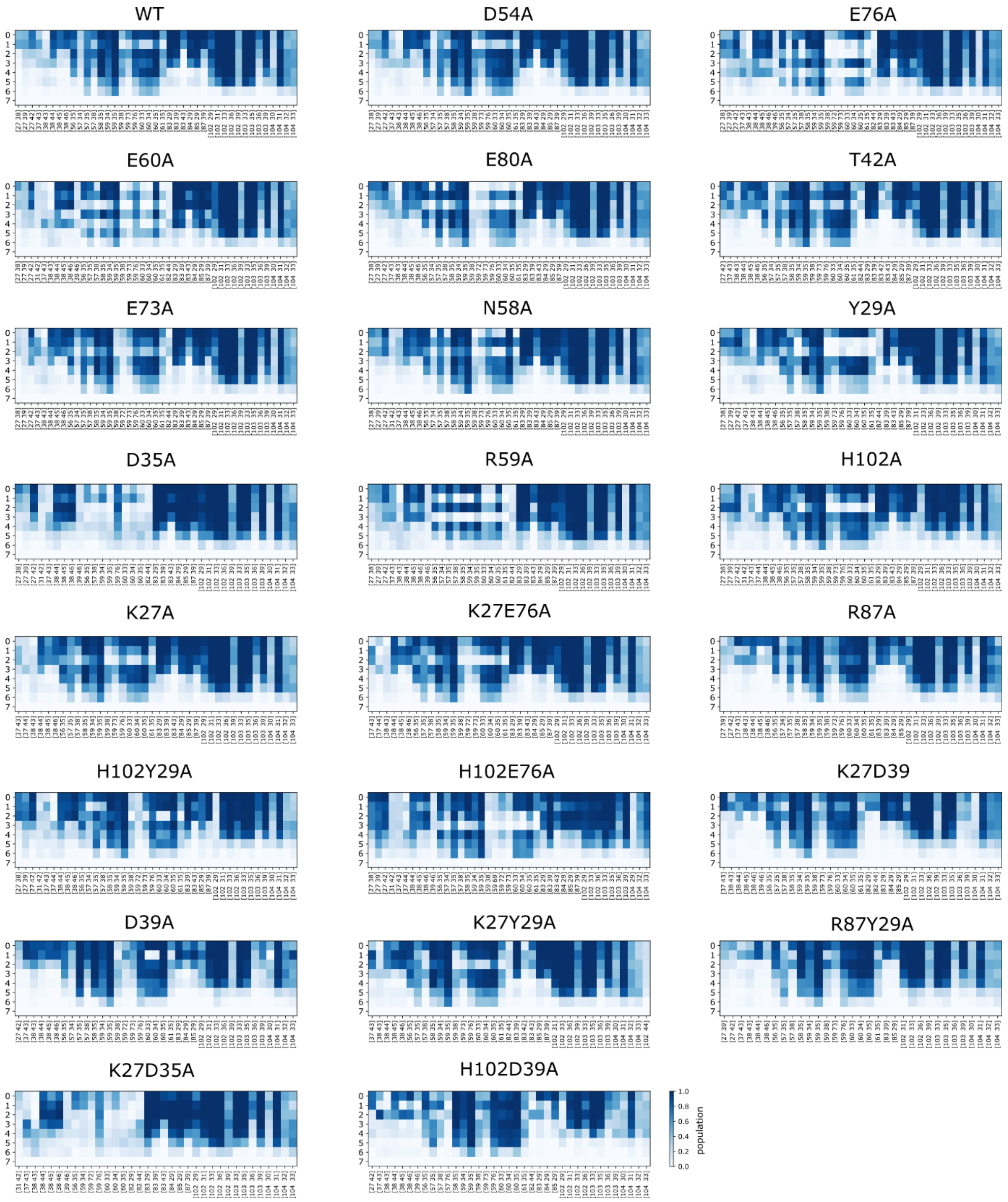


**Figure S4. Interaction fingerprints for the RAMD dissociation trajectories of the wild-type and mutant barnase-barstar complexes.** The interaction fingerprints were computed for the RAMD trajectory clusters resulting from the k-means clustering for WT and mutant Bn-Bs complexes. Clusters are labeled from 0 to 7 (rows). The pairs of residue contacts characterizing each cluster are shown on the x-axis (the first residue index refers to barnase and the second refers to barstar). The population of each pair of residue contacts (columns) is shown with a color scale from blue (highest) to white (lowest).

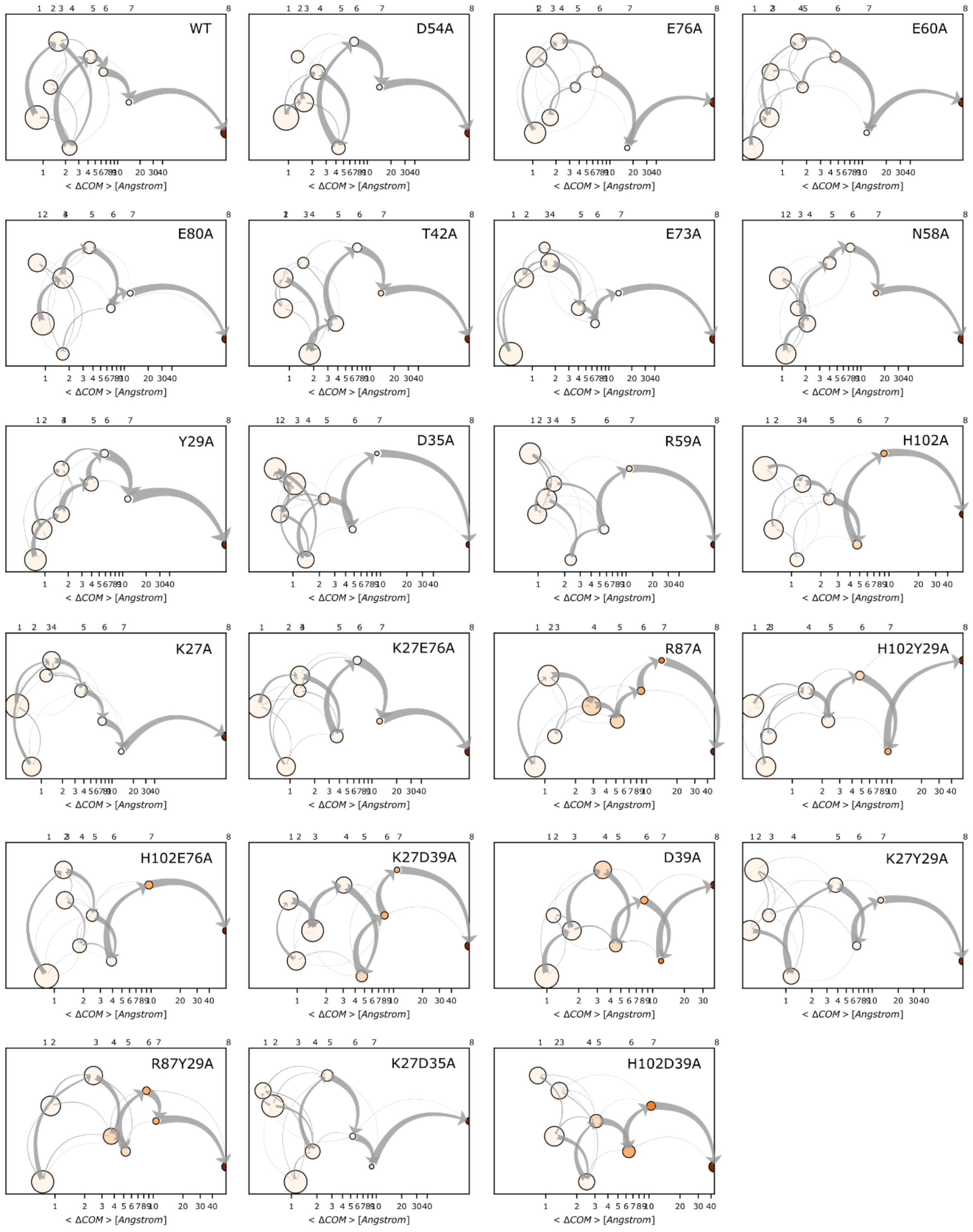


**Figure S5. Schematic representation of the clusters identified during RAMD dissociation trajectories for WT and mutant Bn-Bs complexes**. Each cluster is shown by a node with the size indicating the cluster population (distributed over the y-axis with the corresponding cluster number shown at the top of each plot. Nodes are positioned on a logarithmic scale of increasing mean COM-COM distance between the proteins (x-axis) and the node color denotes the average protein RMSD in the cluster from the starting structure. The gray arrows indicate the total flow between two nodes and their width increases with the number of trajectories having the corresponding transition.

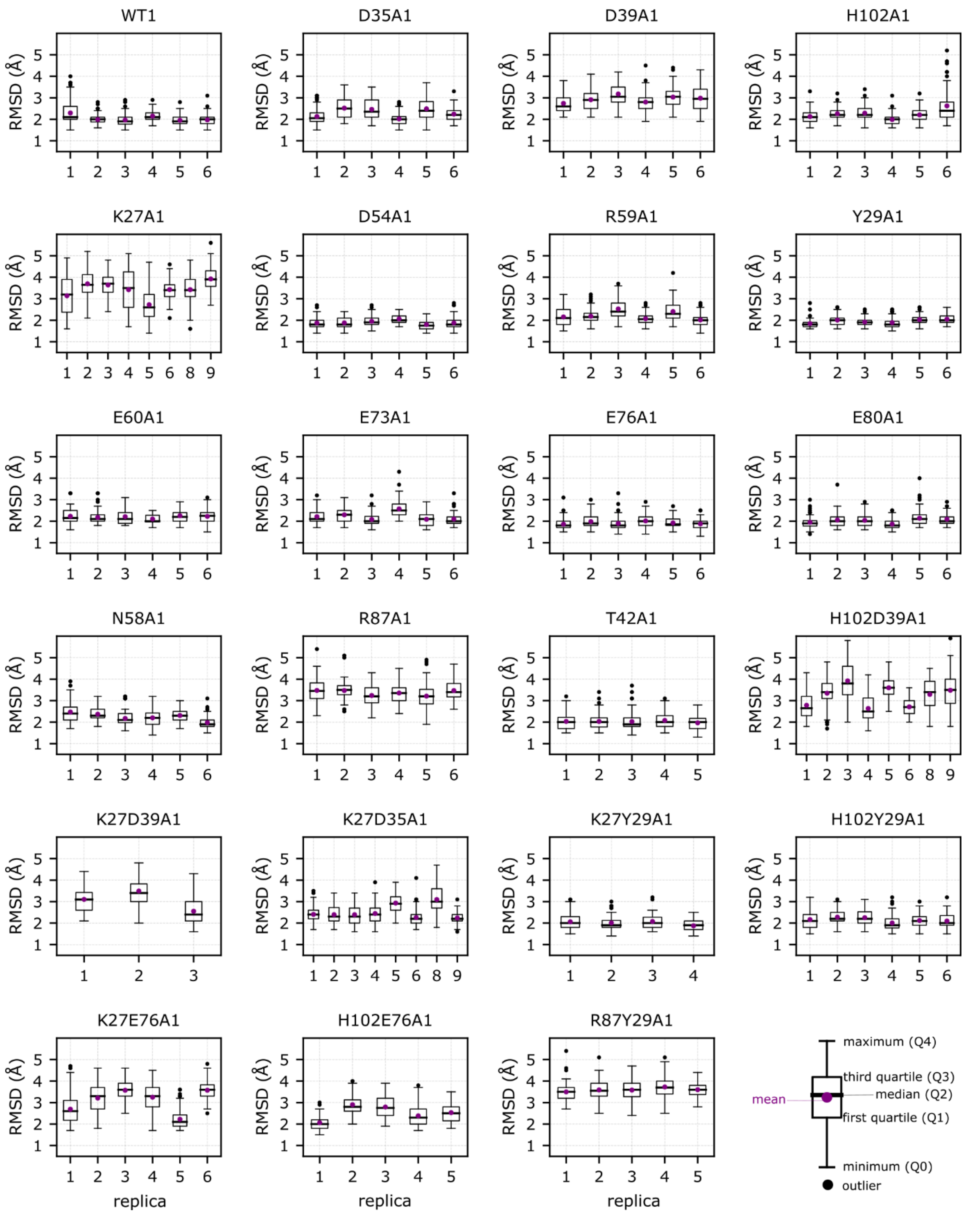


**Figure S6. Root mean squared deviation (RMSD) of Bs after alignment to Bn in the equilibration trajectories for the WT and mutant Bn-Bs complexes.** The RMSD values are less than 6 Å. For the most stable complexes, the mean RMSD is about 2-2.5 Å and similar for all replicas. It is higher and more variable amongst the replicas for the most flexible mutants. RMSDs are shown for 3-6 replica trajectories with mean (purple circle), median (black solid line), first and third quartile (extremes of the box), and extremes of the whiskers at Q3 +

1.5*IQR and Q1 – 1.5*IQR (IQR is interquartile range). Outliers are indicated by black dots. The mean value and standard deviation are indicated above the plot.

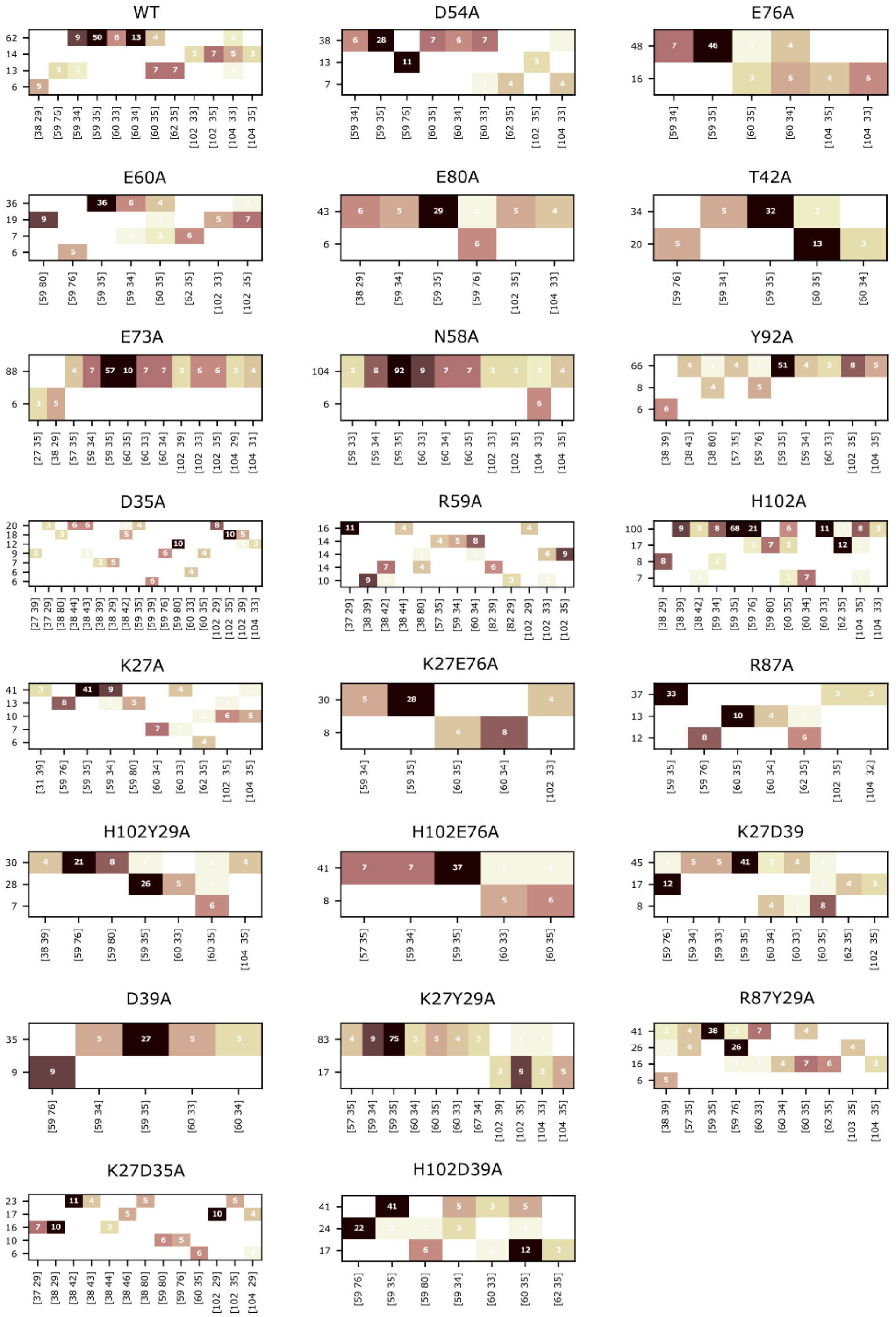


**Figure S7. Reside-residue contacts of the RAMD trajectory clusters for WT and mutant Bn-Bs complexes resulting from the hierarchical clustering of the pre-dissociation frames with fewer than 3 contacts.** Trajectories having the same IFP content are clustered together. Clusters with more than 5 trajectories are shown (1 per row) and ordered on the y-axis from the most populated at the top to the lowest populated at the bottom and labeled by the corresponding number of trajectories. The pairs of residue contacts characterizing the clusters are shown on the x-axis (the first residue index refers to barnase and the second refers to barstar). The occurrence of each pair of residue contacts, computed from MD-IFP analysis, across the trajectories is shown by the corresponding number within each cell and with a color scale from black (highest) to white (lowest).

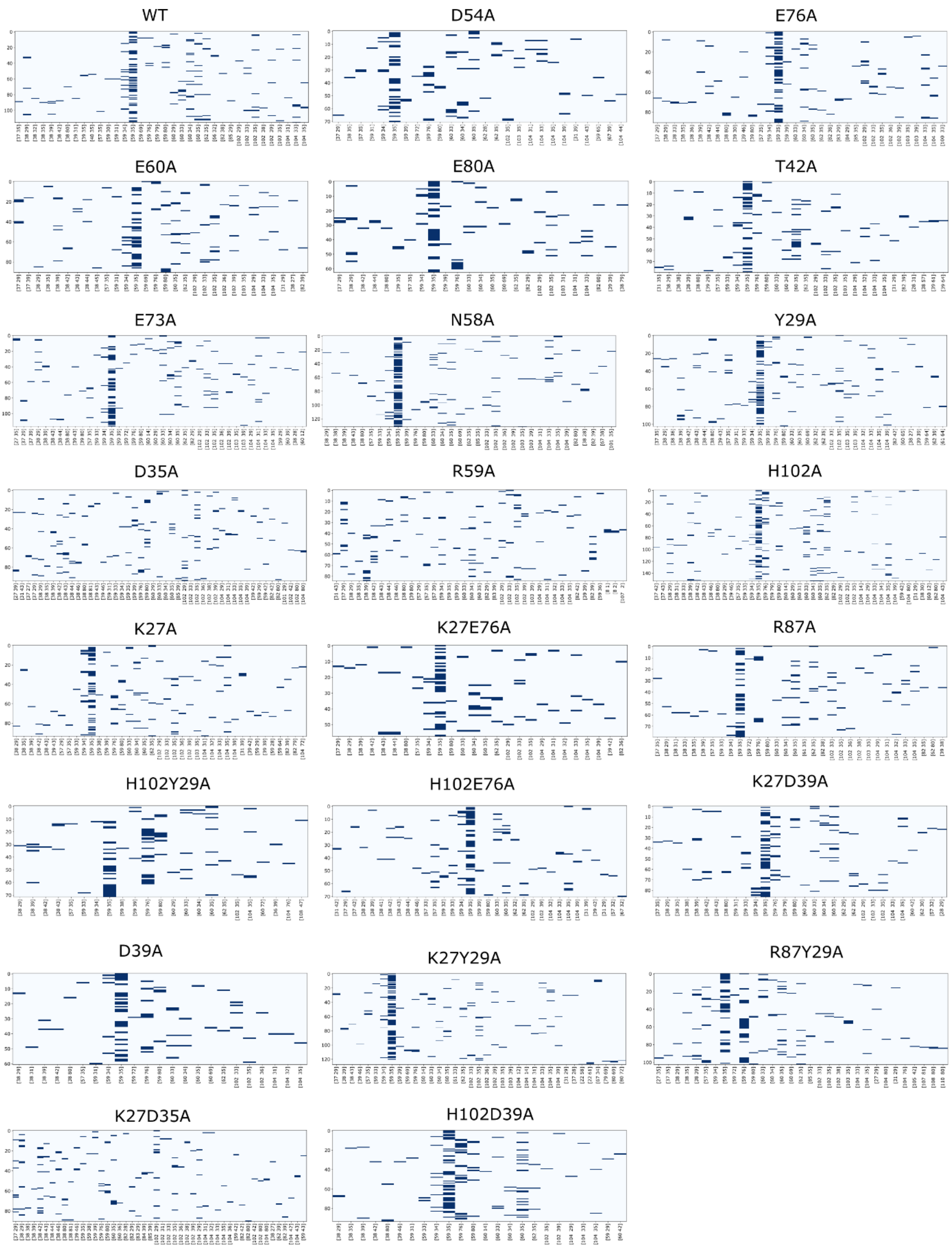


**Figure S8. Residue-residue contacts in the frames from the RAMD trajectories for WT and mutant Bn-Bs complexes with fewer than 3 contacts (excluding the dissociation frames).** The frames are given along the y-axis and the contact pairs on the x-axis (the first residue index refers to barnase and the second refers to barstar).

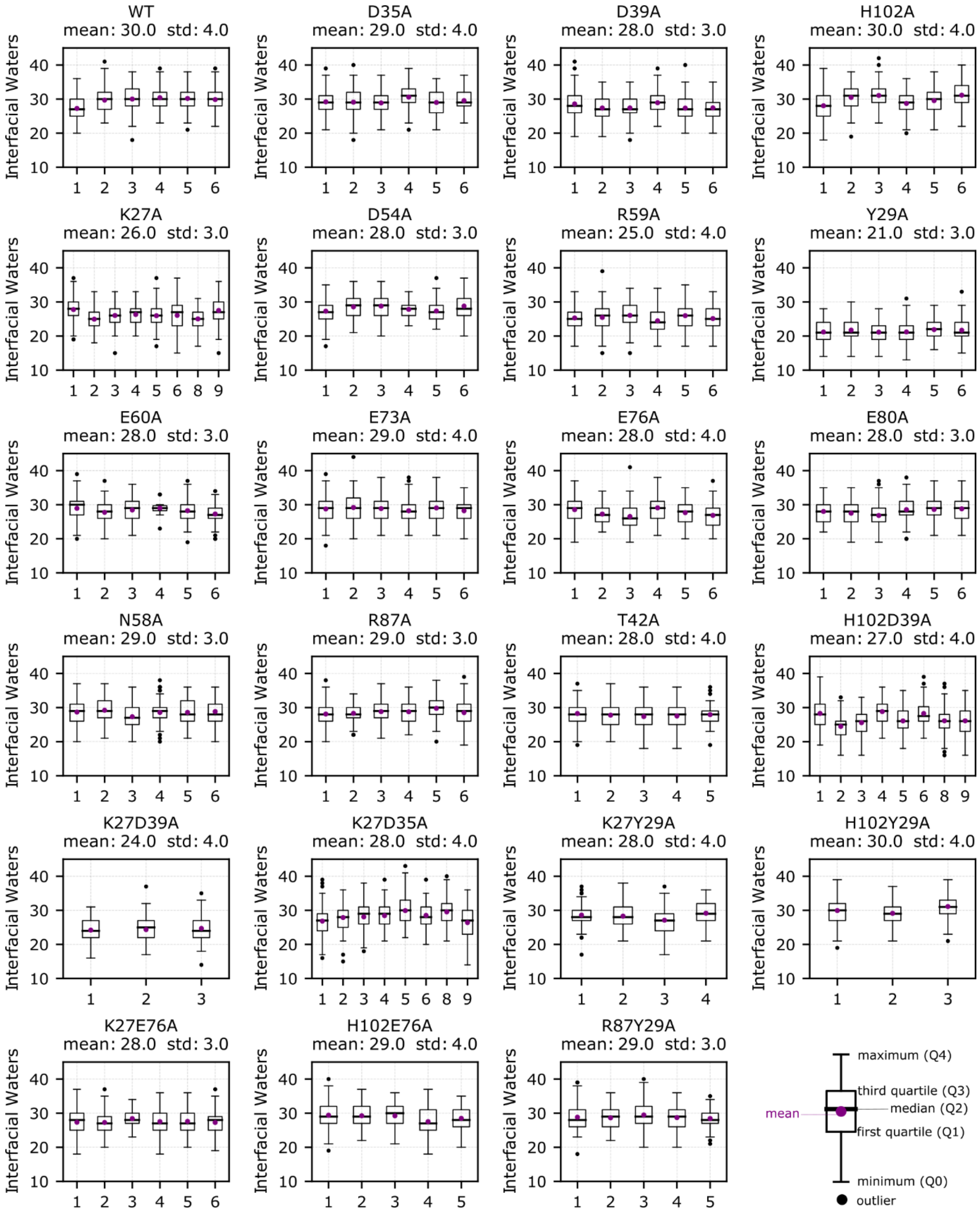


**Figure S9. Number of interfacial waters during equilibration of the WT and mutant Bn-Bs complexes** (see Methods). The numbers of water molecules are similar in all replicas and are shown for 3-6 replica trajectories with mean (purple circle), median (black solid line), first and third quartile (extremes of the box), and extremes of the whiskers at Q3 + 1.5*IQR and Q1 – 1.5*IQR (IQR is interquartile range). Outliers are indicated by black dots. The mean value and standard deviation are indicated above the plot.

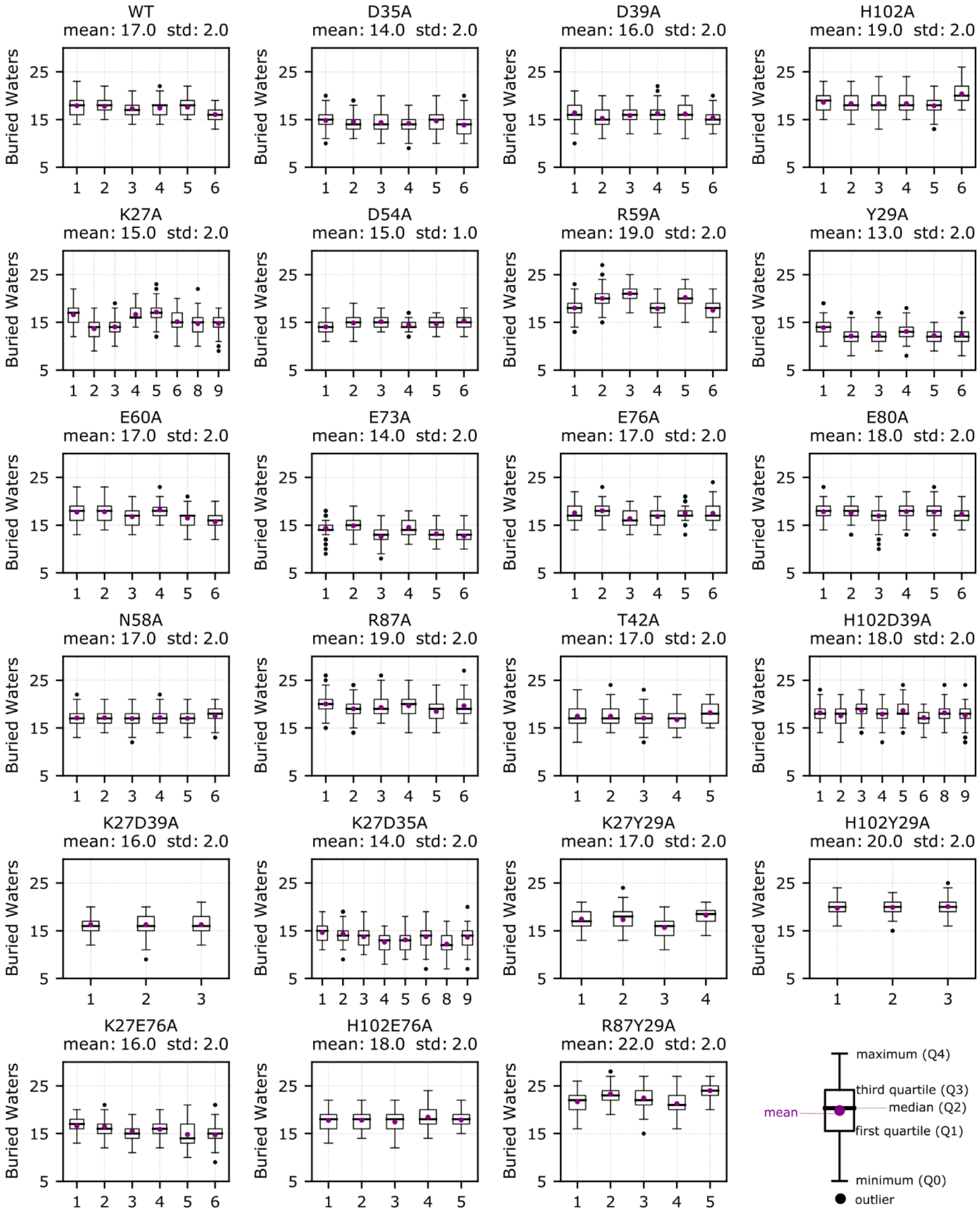


**Figure S10. Number of buried waters during equilibration of the WT and mutant Bn-Bs complexes** (see Methods). The numbers of water molecules are similar in all replicas and are shown for 3-6 replica trajectories with mean (purple circle), median (black solid line), first and third quartile (extremes of the box), and extremes of the whiskers at Q3 + 1.5*IQR and Q1 – 1.5*IQR (IQR is interquartile range). Outliers are indicated by black dots. The mean value and standard deviation are indicated above the plot.

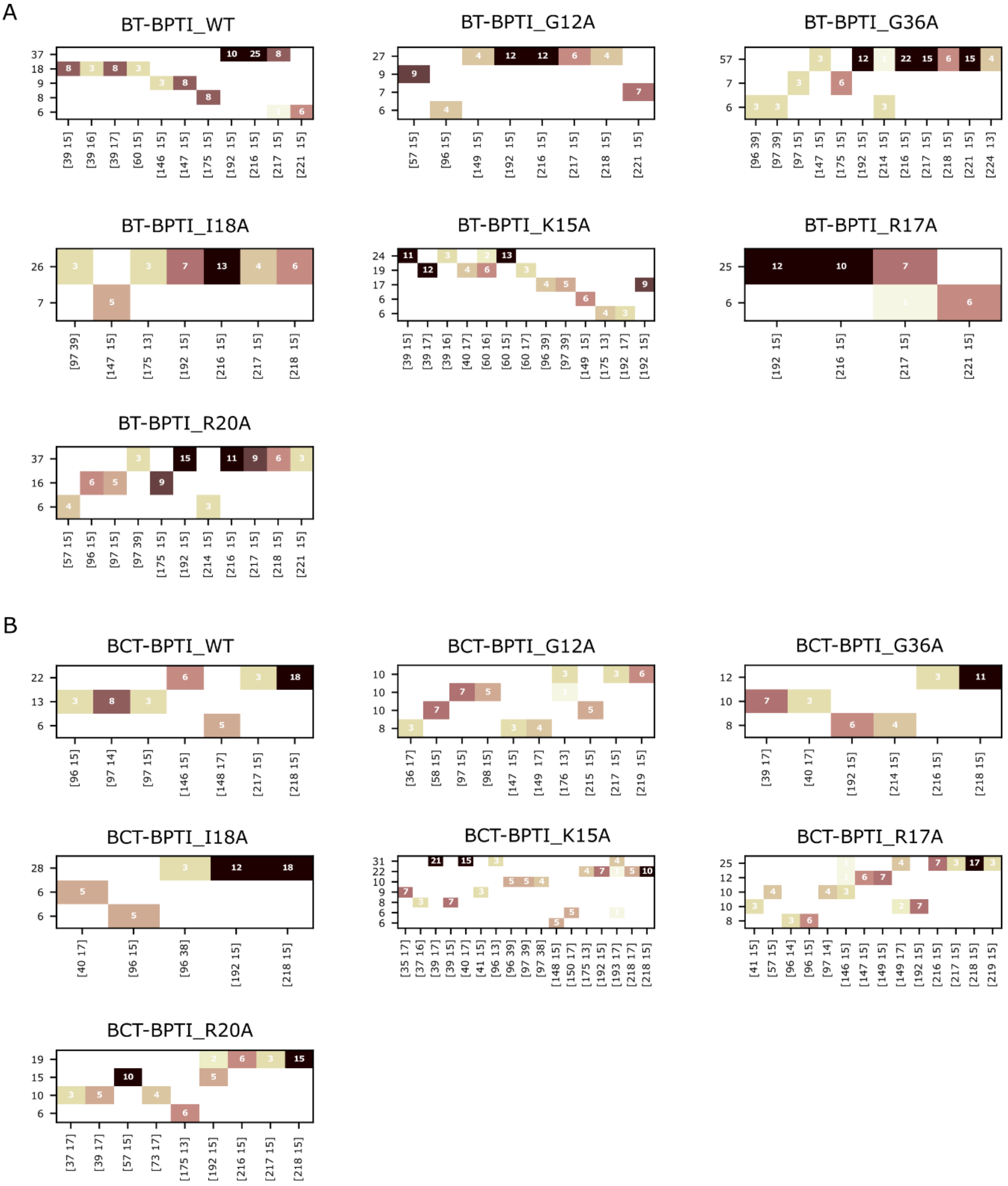


**Figure S12. Reside-residue contacts of the RAMD trajectory clusters for WT and mutant BT-BPTI and BCT-BPTI complexes resulting from the hierarchical clustering of the pre-dissociation frames with fewer than 3 contacts.** Trajectories having the same IFP content are clustered. Clusters with more than 5 trajectories are shown and ordered on the y-axis from the most populated to the lowest populated, and labeled by the corresponding number of trajectories. The pairs of residue contacts characterizing the clusters are shown on the x-axis (the first residue index refers to BT/BCT and the second refers to BPTI). The occurrence of each pair of residue contacts, computed from MD-IFP analysis, across the trajectories is shown by the corresponding number within each cell and with a color scale from black (highest) to white (lowest).

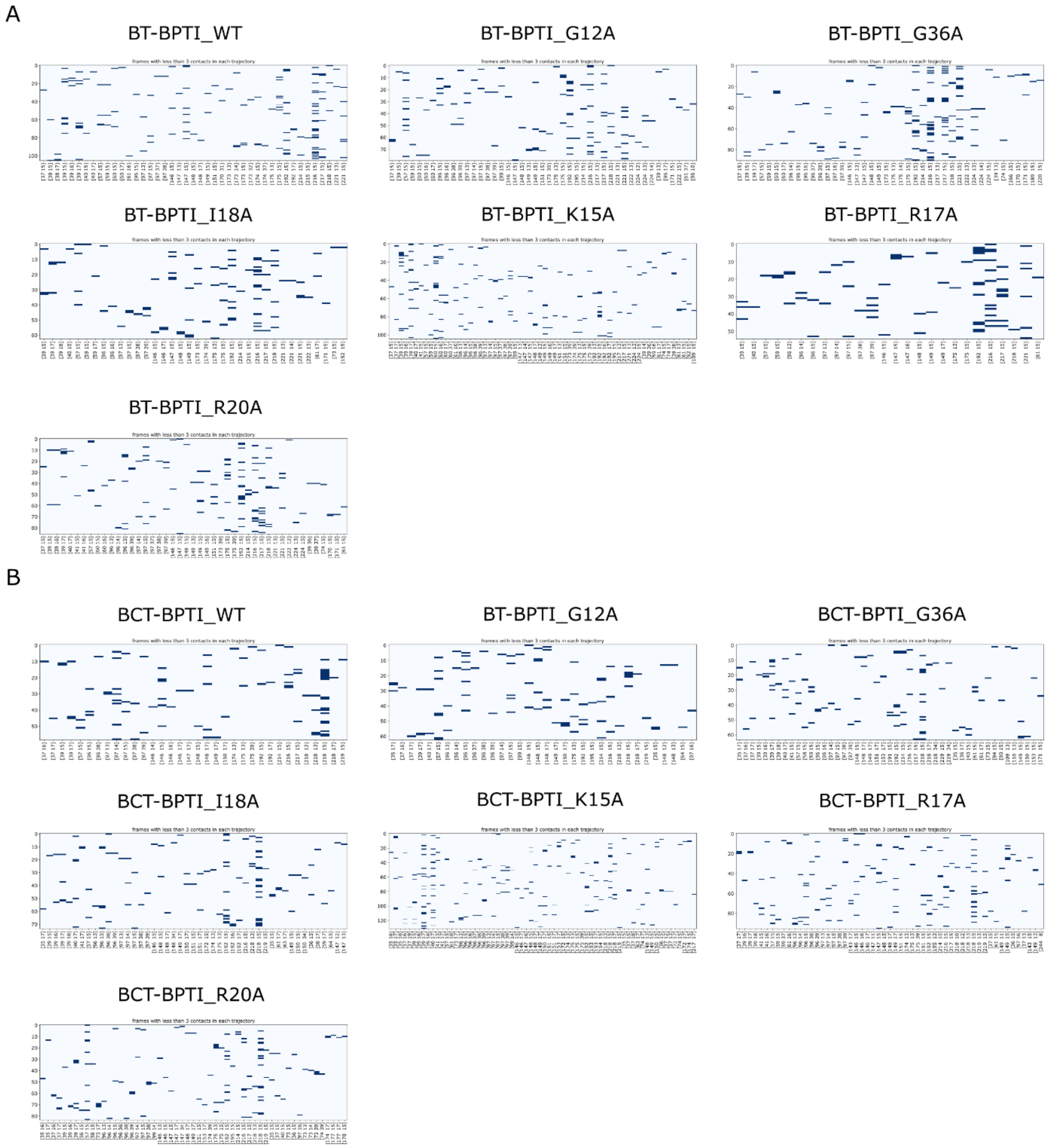


**Figure S13. Residue-residue contacts in the frames from the RAMD trajectories with fewer than 3 contacts (excluding the dissociation frames) for A) BT-BPTI and B) BCT-BPTI WT and mutant complexes.** The frames are given along the y-axis and the contact pairs on the x-axis (the first residue index refers to BT/BCT and the second refers to BPTI).

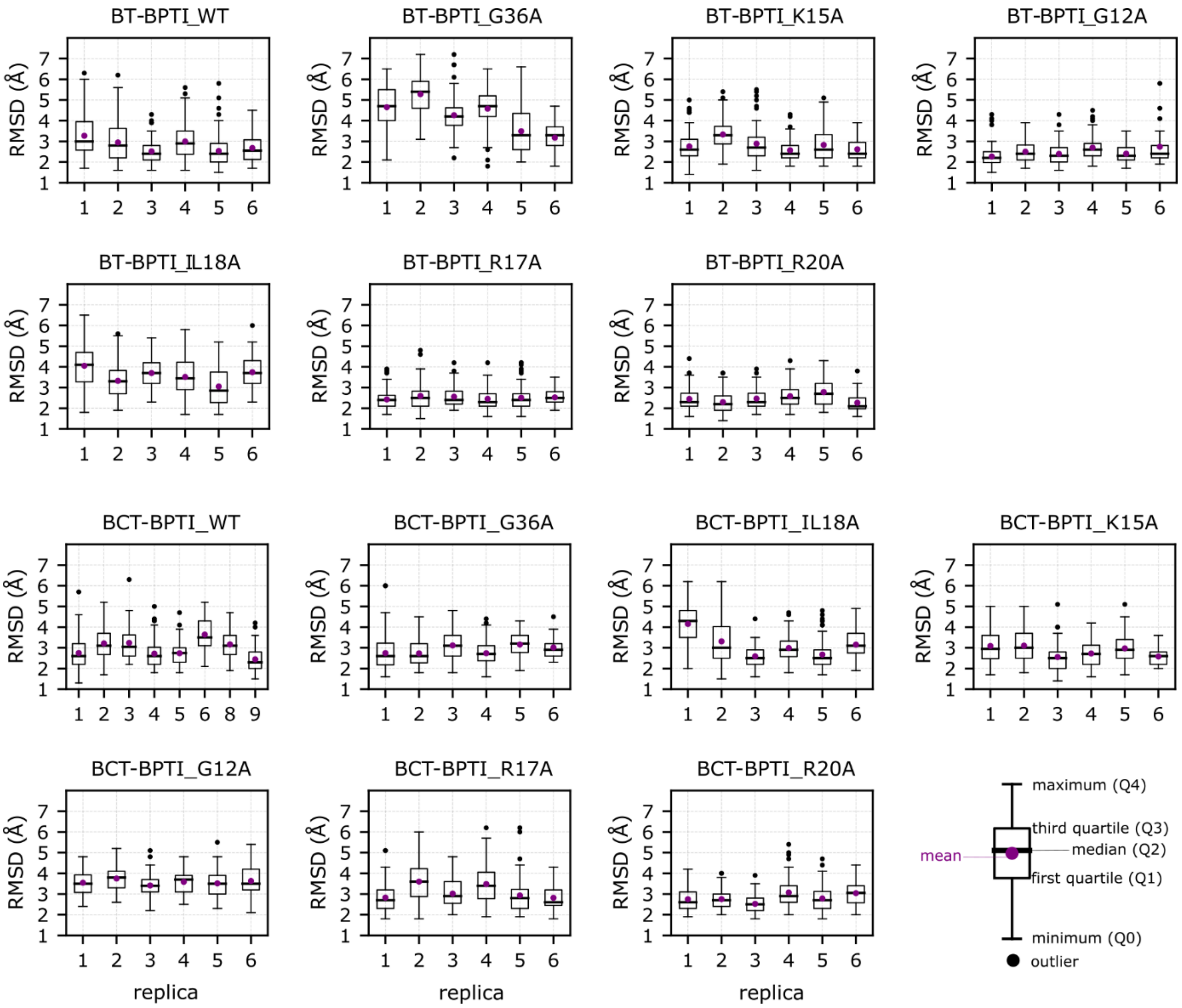


**Figure S14. RMSD of BPTI after alignment to BT/BCT in the equilibration trajectories for the BT-BPTI and CBT-BPTI WT and mutant complexes**. RMSD values are less than 7 Å, with a mean of about 2.5-3 Å for the most stable BT/BCT-BPTI complexes and higher for the most flexible mutants. RMSDs are shown for 6 replica trajectories with mean (purple circle), median (black solid line), first and third quartile (extremes of the box), and extremes of the whiskers at Q3 + 1.5*IQR and Q1 – 1.5*IQR (IQR is interquartile range). Outliers are indicated by black dots.

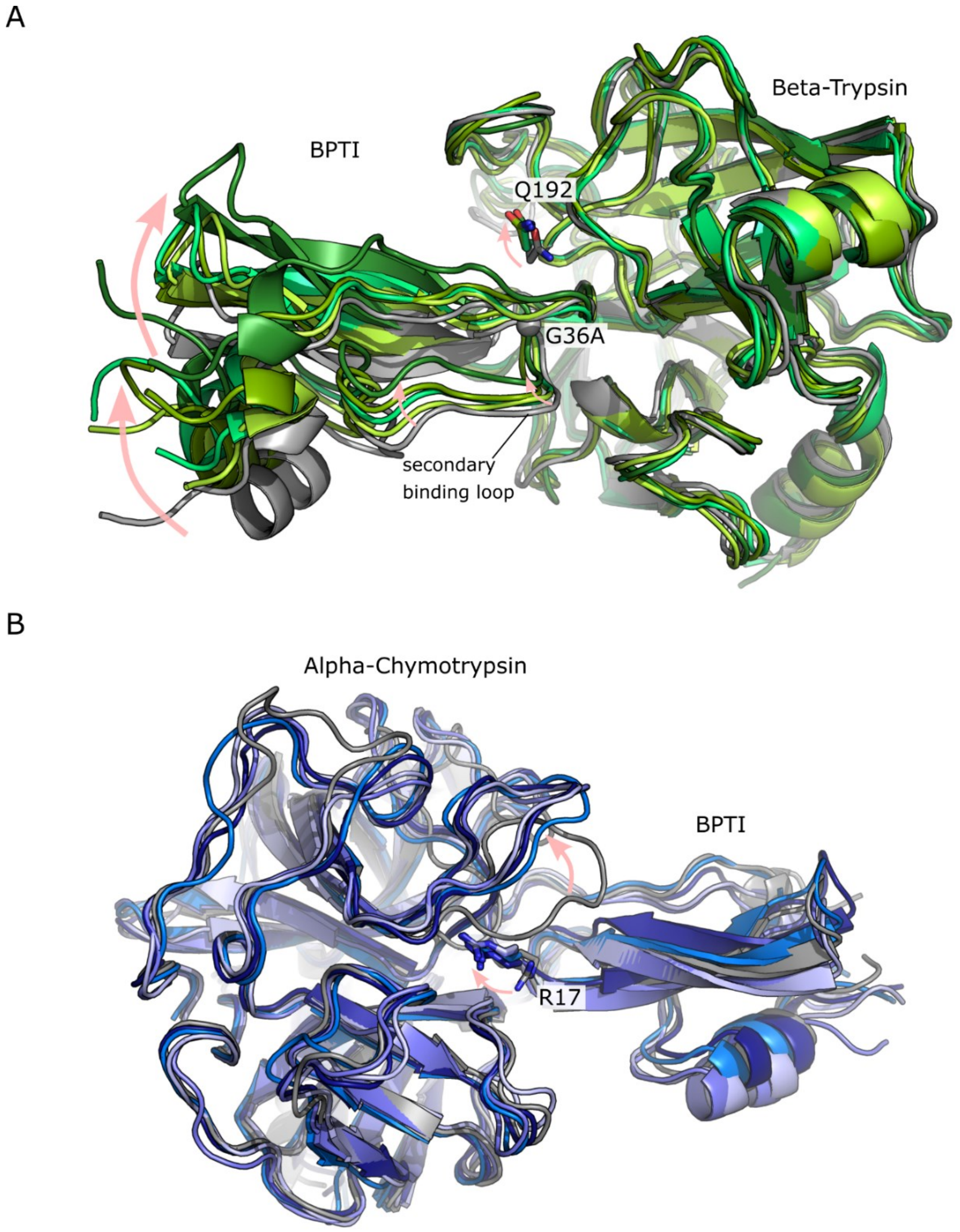


**Figure S15. Comparison of the structures of WT and selected mutant complexes with BPTI.** **A)** Superposition of the WT BT-BPTI complex (grey cartoon) and four replicas of the G36A BT-BPTI complex (in different shades of the green cartoon) taken from the equilibration trajectories. The G36A point mutation is shown as a grey sphere and labeled. The shifts in the BPTI structure with respect to the WT complex are highlighted by arrows, as well as the shift in the orientation of Q192 of BT. **B)** Superposition of the WT BCT-BPTI complex (grey cartoon) and four replicas of the K15A BCT-BPTI complex (in different shades of the blue cartoon), taken from the equilibration trajectories. The shifts in the loop composing the S2’ pocket in BCT, and the shift in the orientation of R17 of BPTI are highlighted by arrows.

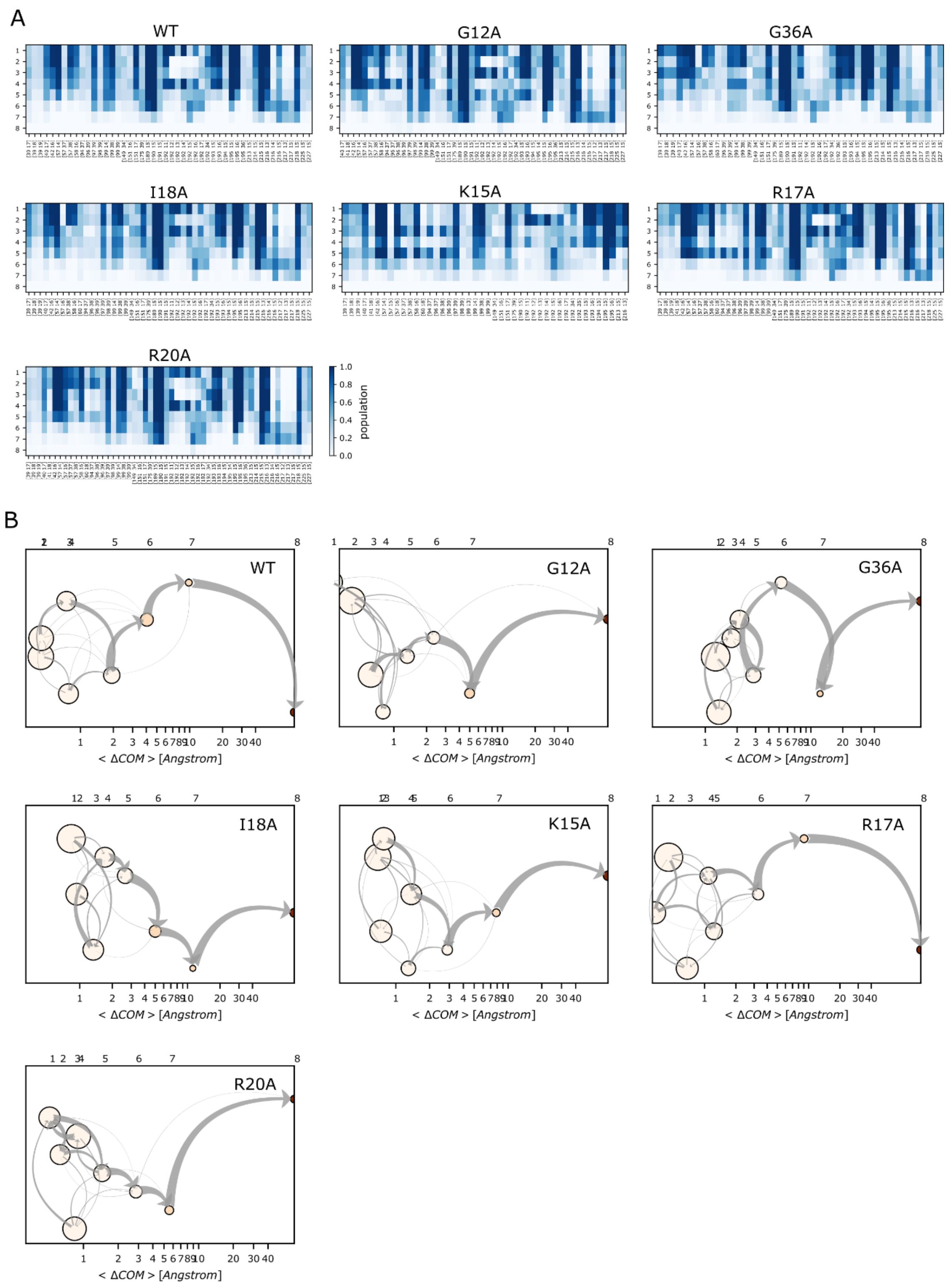


**Figure S16**. **Analysis of RAMD trajectories for the BT-BPTI WT and mutant complexes. A)** IFP composition of the trajectory clusters resulting from the k-means clustering. Clusters are labeled from 1 to 8 (rows). The population of each pair of residue contacts is shown with a color scale from blue (highest) to white (lowest). **B)** Schematic representation of the clusters visited during the RAMD dissociation trajectories. Clusters are labeled above the plots and are ordered by increasing mean COM-COM distance between proteins (x-axis). Cluster color indicates the average protein RMSD in the cluster from the starting structure.

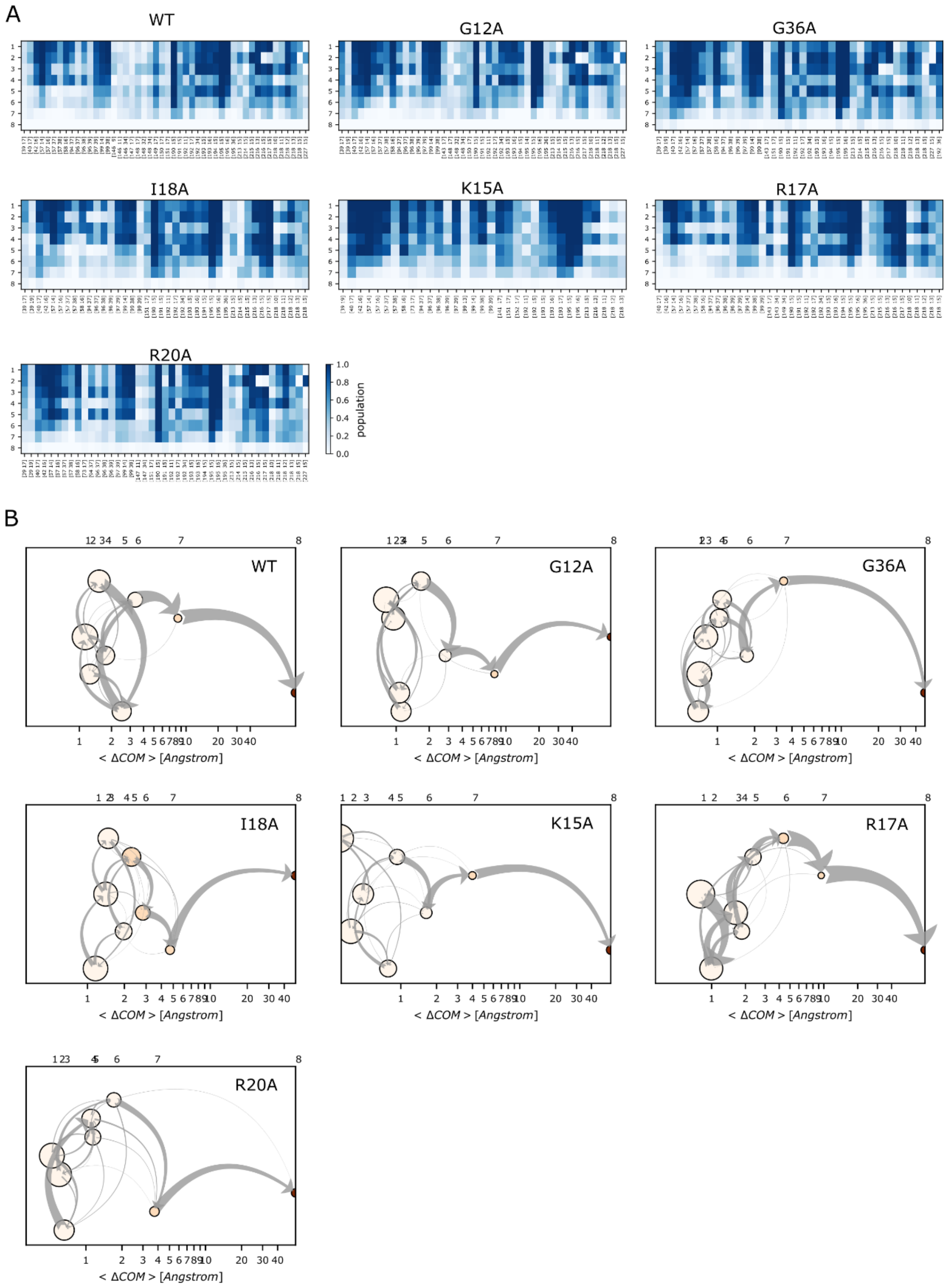


**Figure S17**. **Analysis of RAMD trajectories for the BCT-BPTI WT and mutant complexes. A)** IFP composition of the trajectory clusters resulting from the k-means clustering. Clusters are labeled from 1 to 8 (rows). The population of each pair of residue contacts is shown with a color scale from blue (highest) to white (lowest). **B)** Schematic representation of the clusters visited during the RAMD dissociation trajectories. Clusters are labeled above the plots and are ordered by increasing mean COM-COM distance between proteins (x-axis). Cluster color indicates the averaged protein RMSD in the cluster from the starting structure.

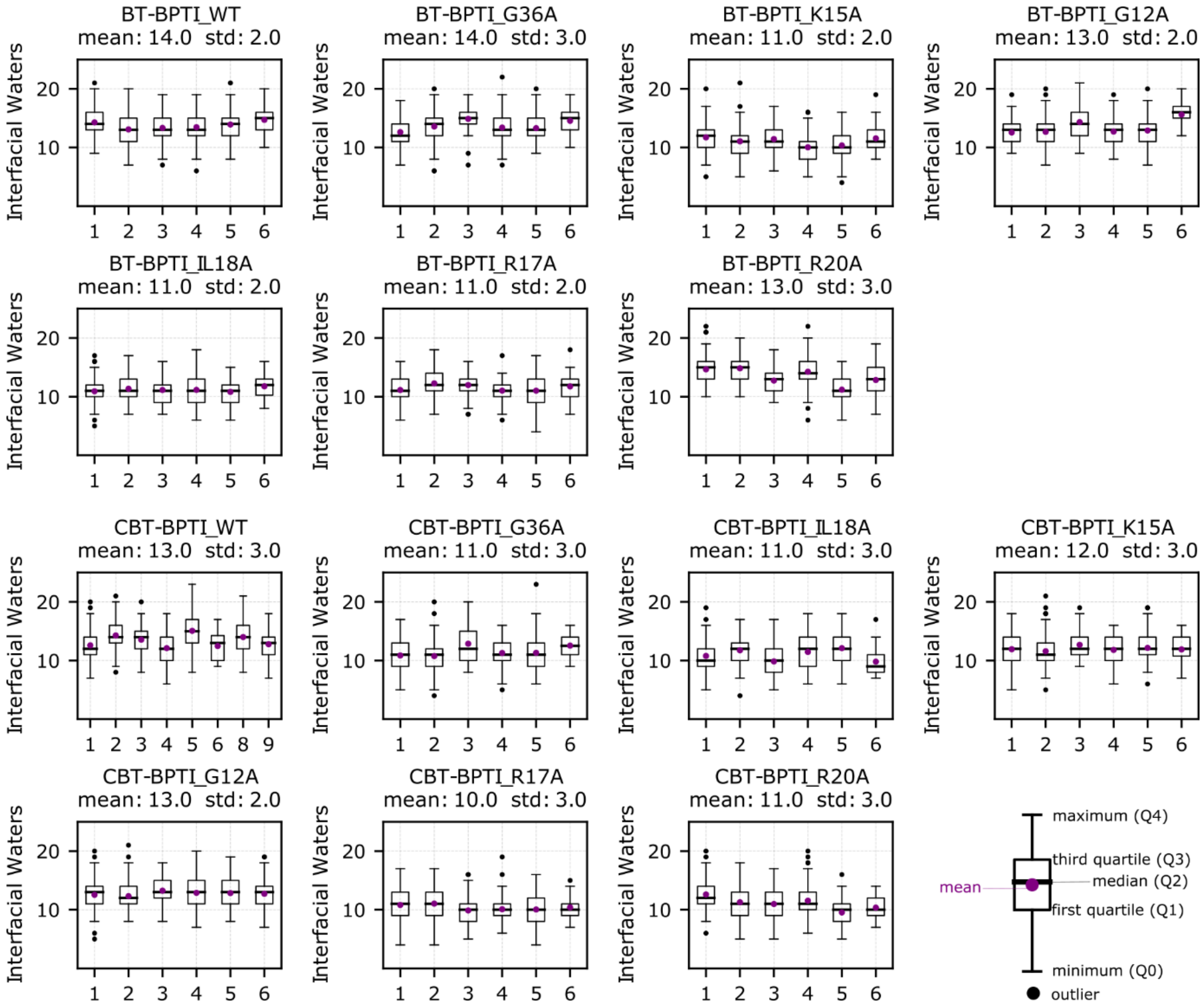


**Figure S11. Number of interfacial waters during equilibration of the WT and mutant BT-BPTI and BCT-BPTI complexes** (see Methods). The numbers of water molecules are similar in all replicas and are shown for 3-6 replica trajectories with mean (purple circle), median (black solid line), first and third quartile (extremes of the box), and extremes of the whiskers at Q3 + 1.5*IQR and Q1 – 1.5*IQR (IQR is interquartile range). Outliers are indicated by black dots. The mean value and standard deviation are indicated above the plot.

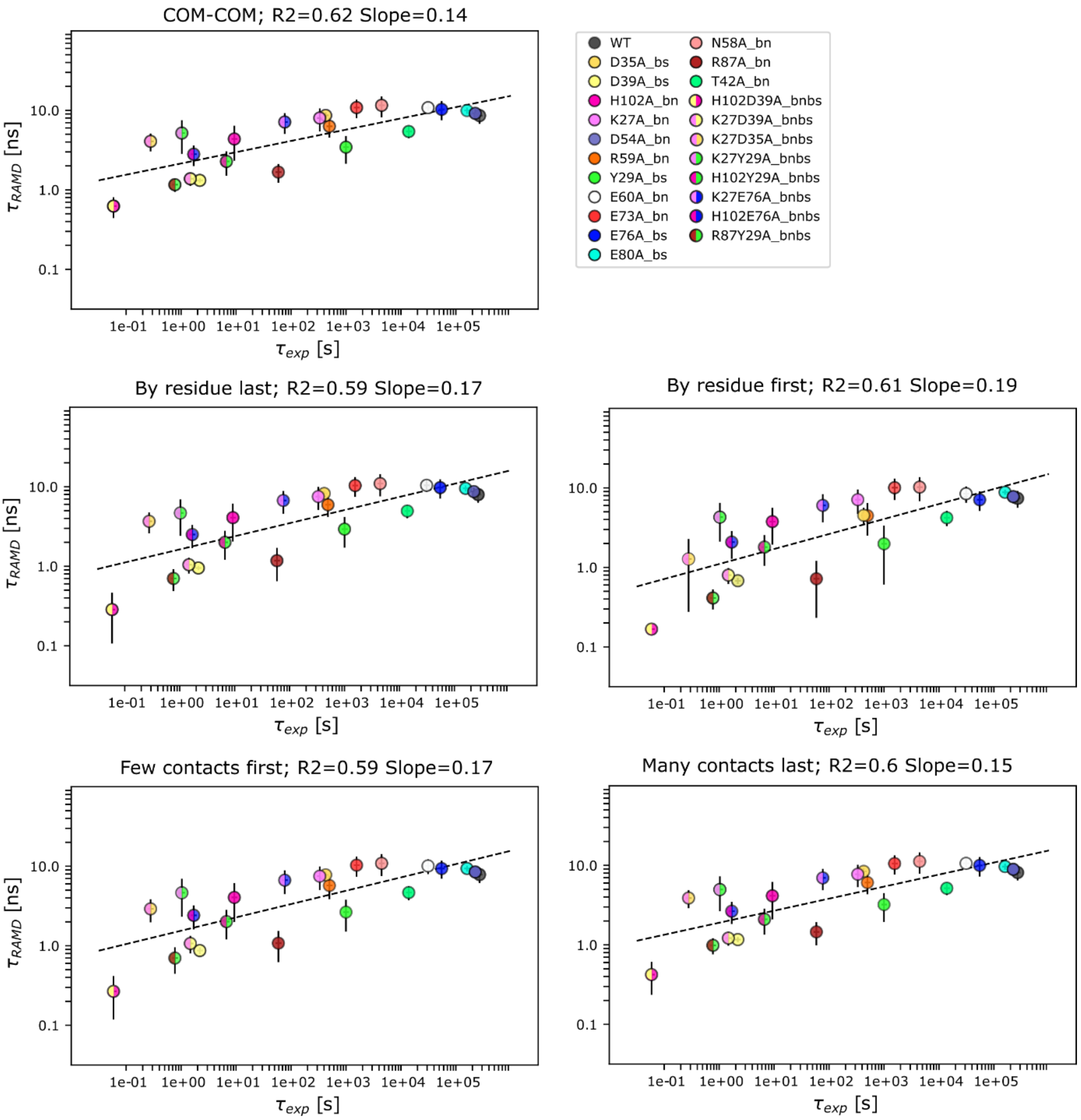


**Figure S18. RAMD residence time, $\tau_{RAMD}$, computed for wild-type and mutant Bn-Bs complexes vs experimental residence time ($\tau_{exp}$, inverse of $k_{off}$).** Results are shown for the different definitions of residence time (see Methods) which all give a similar correlation with an R2 value of about 0.6. A random force magnitude of 19 kcal/mol/Å was used. The error bars indicate computed standard deviations and the dashed line shows the straight-line fit to all the data with the given R2 value and slope.

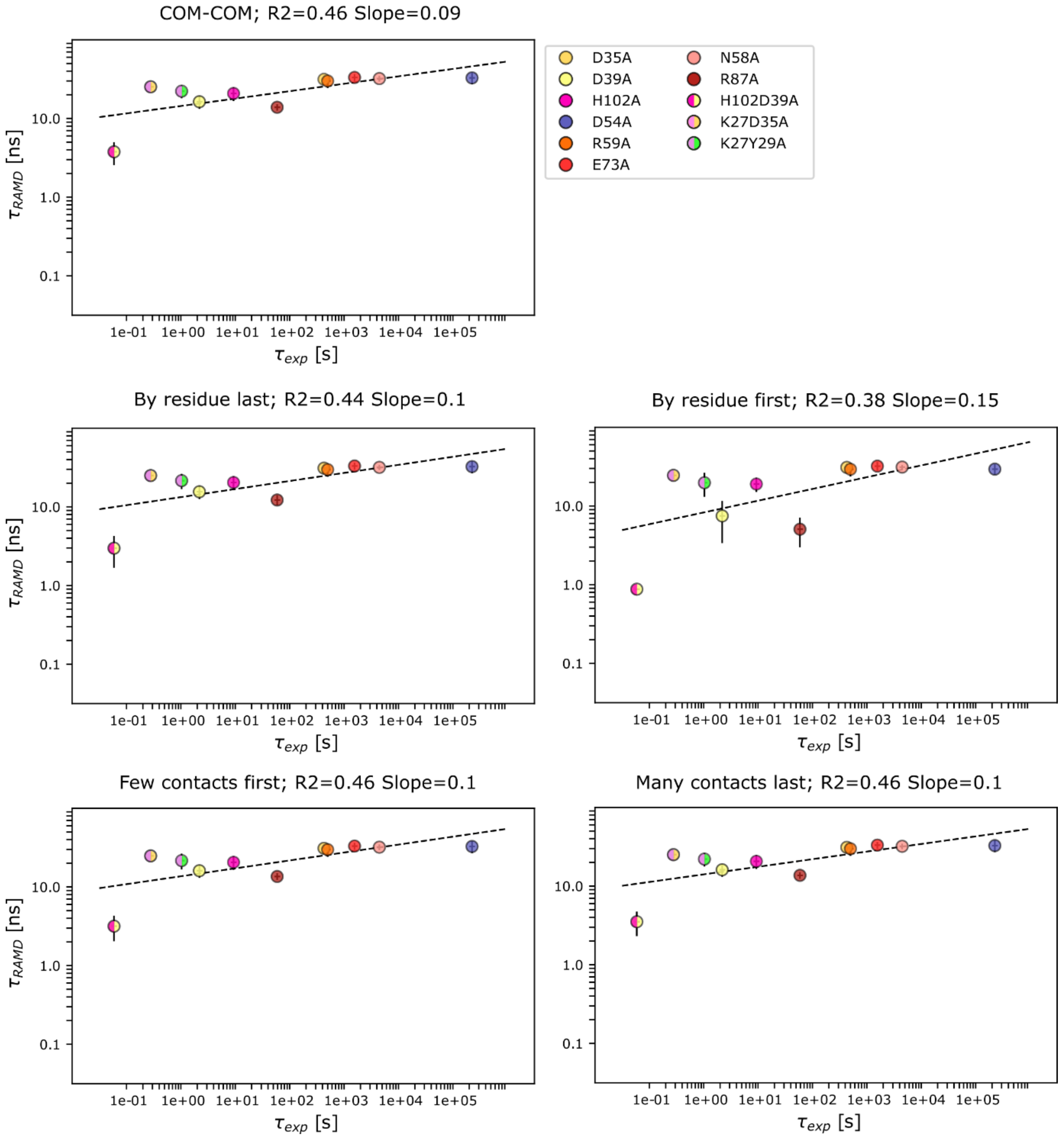


**Figure S19. RAMD residence times for a subset of Bn-Bs mutants computed using a random force magnitude of 17 kcal/mol/Å vs experimental residence time (inverse of $k_{off}$), for the different definitions of residence time** (see Methods). The low random force magnitude results in a low slope and low correlation of computed and experimental residence times. The error bars indicate computed standard deviations and the dashed line shows the straight-line fit to all the data with the given R2 value and slope.

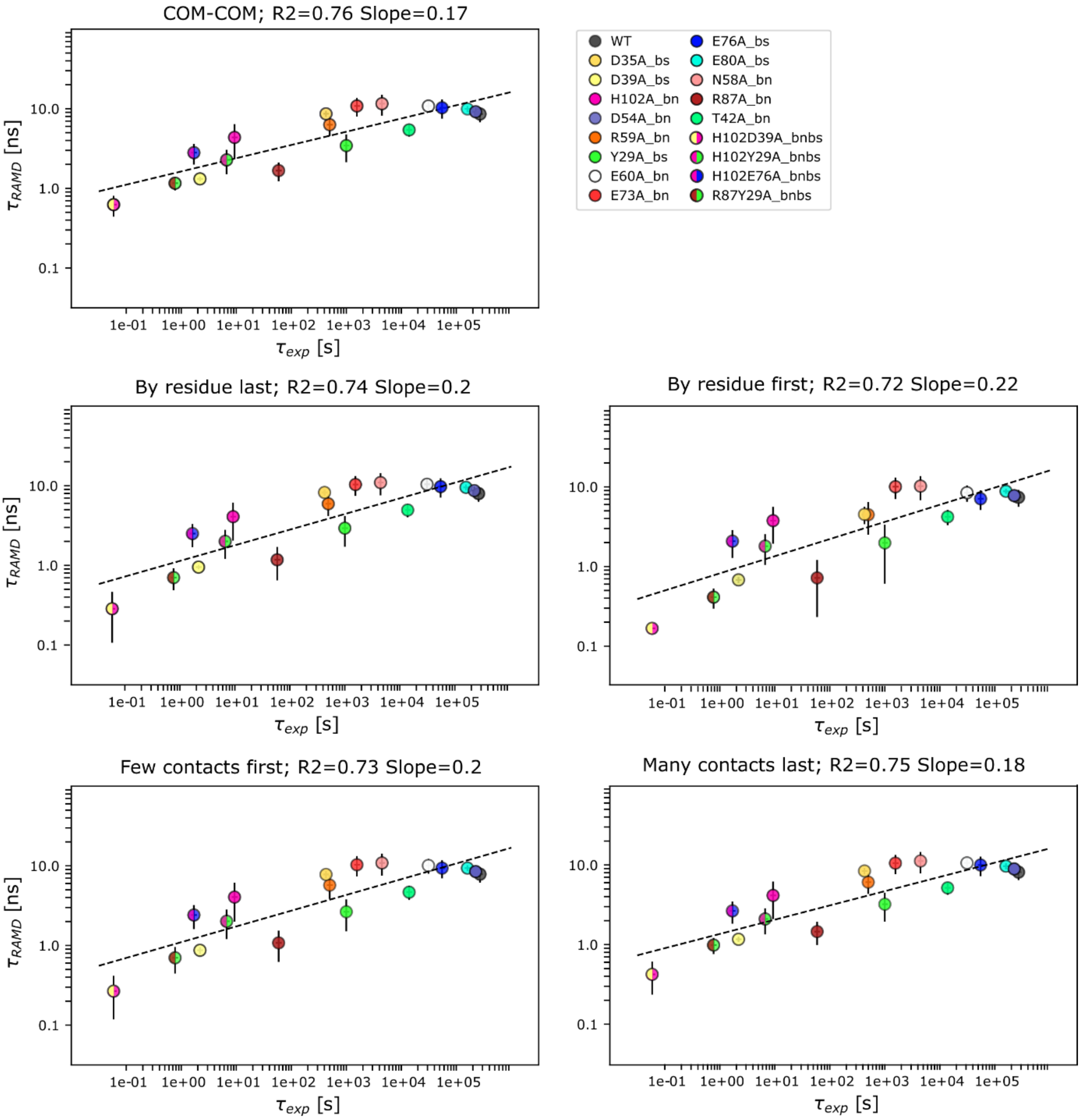


**Figure S20. RAMD residence time for the Bn-Bs complexes computed with a random force magnitude of 19 kcal/mol/Å vs experimental residence times (inverse of $k_{off}$), for the different definitions of residence time (see Methods), without the single and double K27A mutants.** A good correlation with an R2 value of about 0.75 is obtained for all residence time definitions. The error bars indicate computed standard deviations and the dashed line shows the straight-line fit to all the data with the given R2 value and slope.

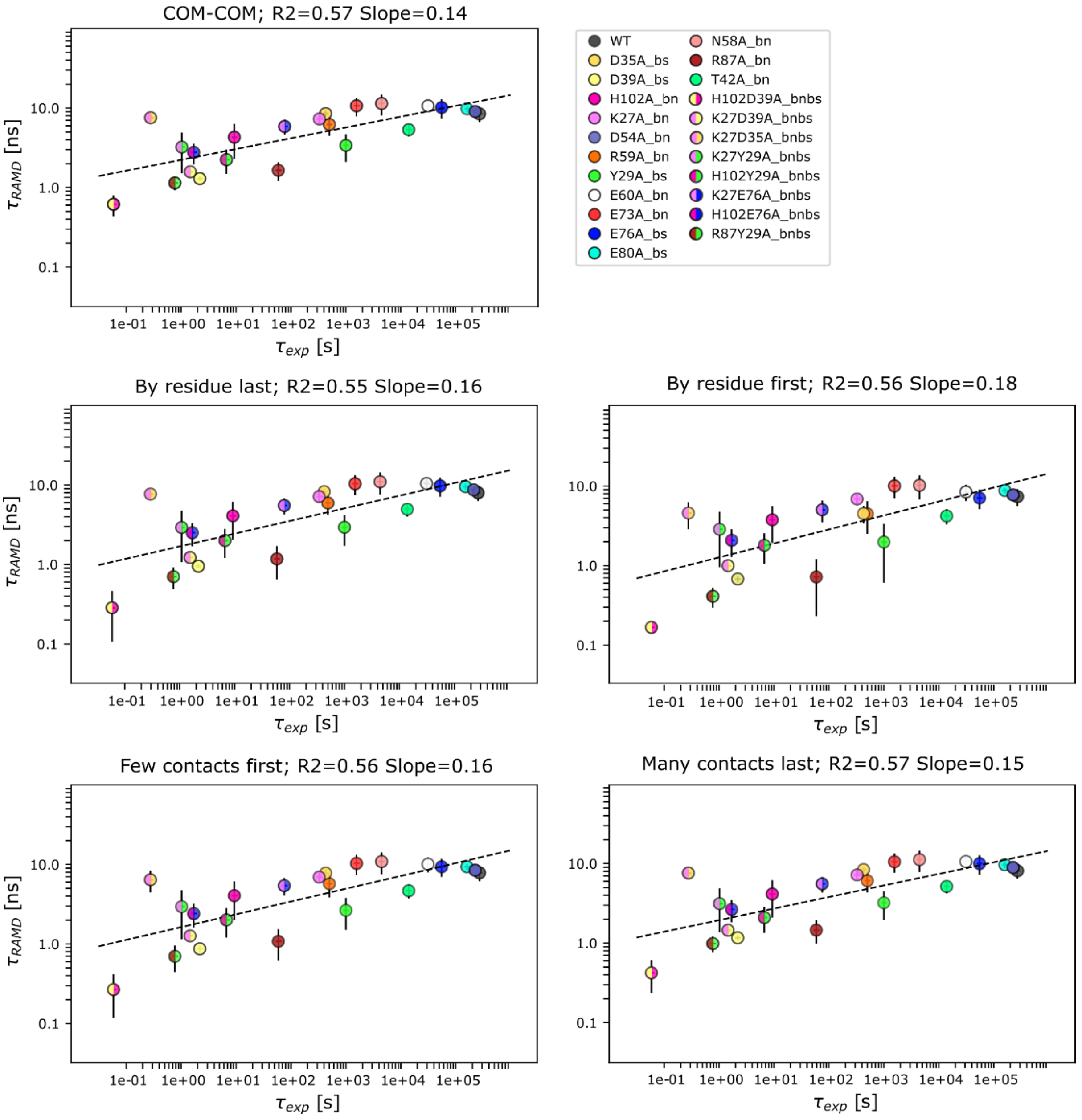


**Figure S21. RAMD residence time for the Bn-Bs complexes computed with a random force magnitude of 19 kcal/mol/Å vs experimental residence times (inverse of $k_{off}$) for the different definitions of residence time (see Methods), with deprotonated E73bn in the single and double K27Abn mutants.** A correlation with an R2 value of about 0.55 is obtained for all residence time definitions. The error bars indicate computed standard deviations and the dashed line shows the straight-line fit to all the data with the given R2 value and slope.

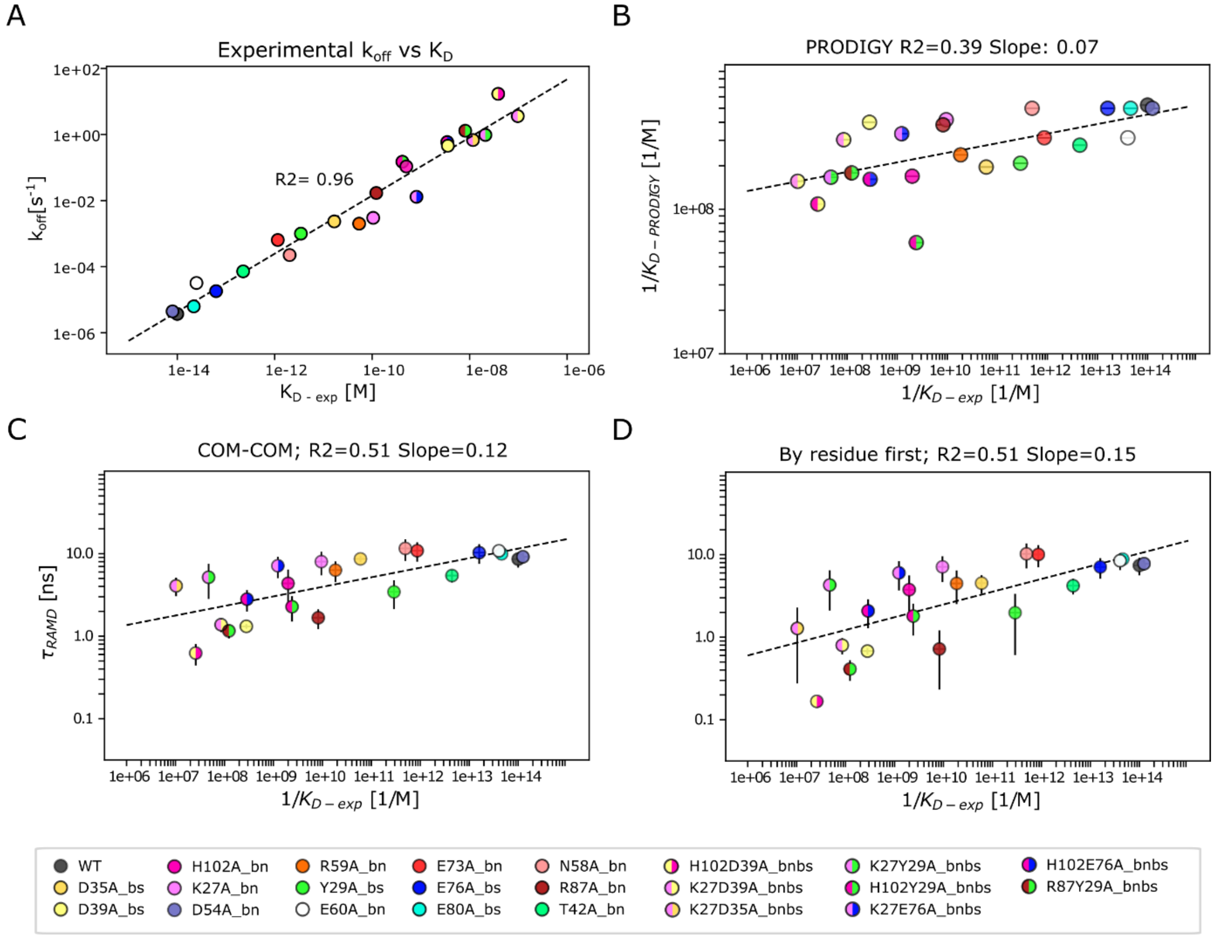


**Figure S22. Comparison with the experimentally measured inverse value of $K_d$ for the Bn-Bs complexes**. **A)** Correlation between experimental normalized $k_{off}$ and $K_d$ values. **B)** PRODIGY prediction of $K_d$ vs experimental values with an R2 value of about 0.4. **C-D)** RAMD residence time computed with a random force magnitude of 19 kcal/mol/Å vs experimental inverse values of $K_d$. Correlations are shown for the ''COM-COM'' and ''By residue first'' definitions of residence time (see Methods) with an R2 of 0.51 for both approaches. The error bars indicate computed standard deviations and the dashed line shows the straight-line fit to all the data with the given R2 value and slope.

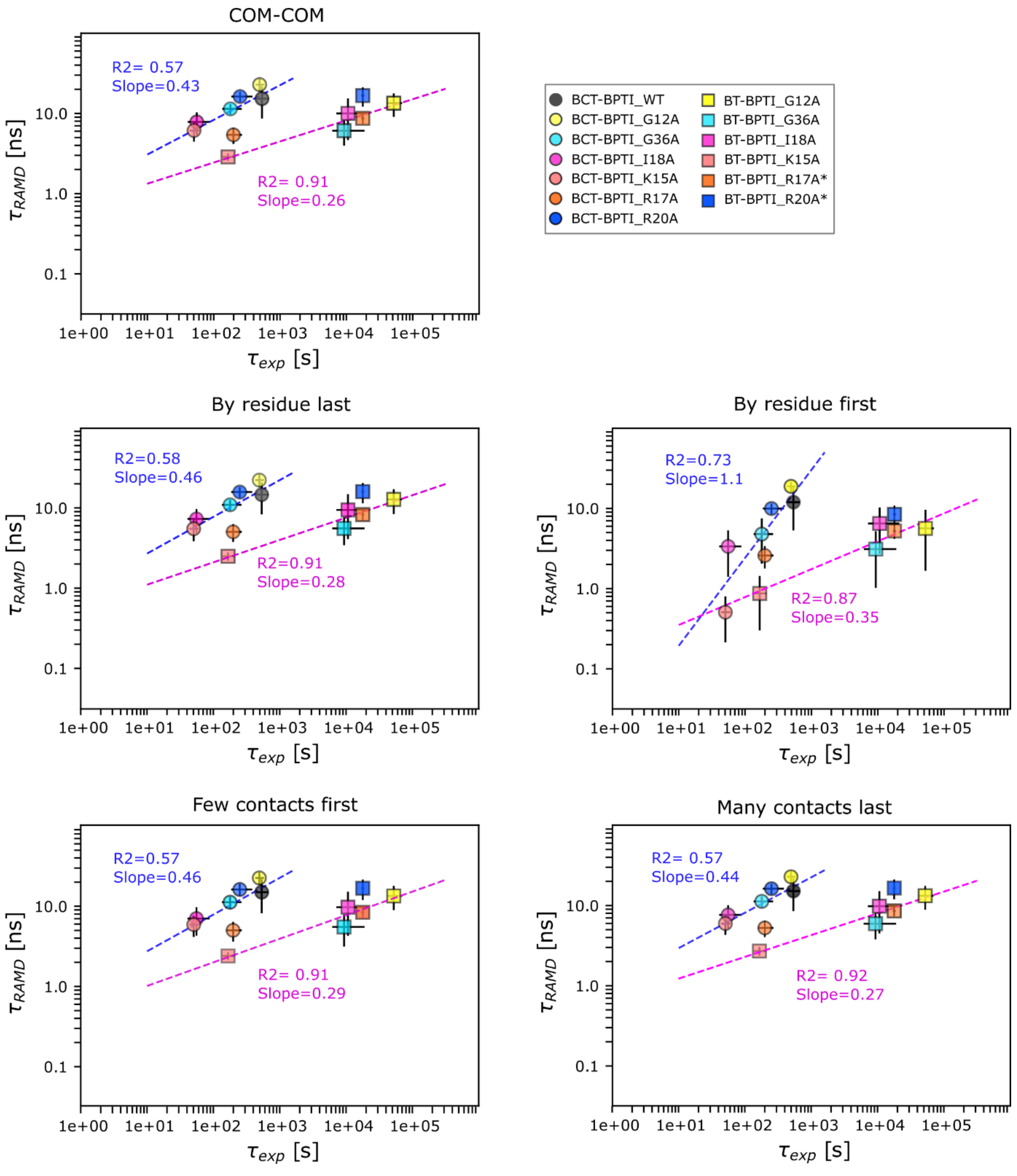


**Figure S23. RAMD residence time, $\tau_{RAMD}$, computed for wild-type and mutant BT-BPTI and BCT-BPTI complexes vs experimental residence time ($\tau_{exp}$, inverse of $k_{off}$).** Results are shown for the different definitions of residence time (see Methods). A random force magnitude of 17 kcal/mol/Å was used. The error bars indicate standard deviations and the dashed lines show straight-line fits, with the given R2 value and slope, in magenta and blue for the BT-BPTI and BCT-BPTI complexes, respectively. BT mutants indicated with * in the legend have experimental $k_{off} < 5.6 \times 10^{-5}$ $s^{-1}$ and were not used for determining the straight line fit.

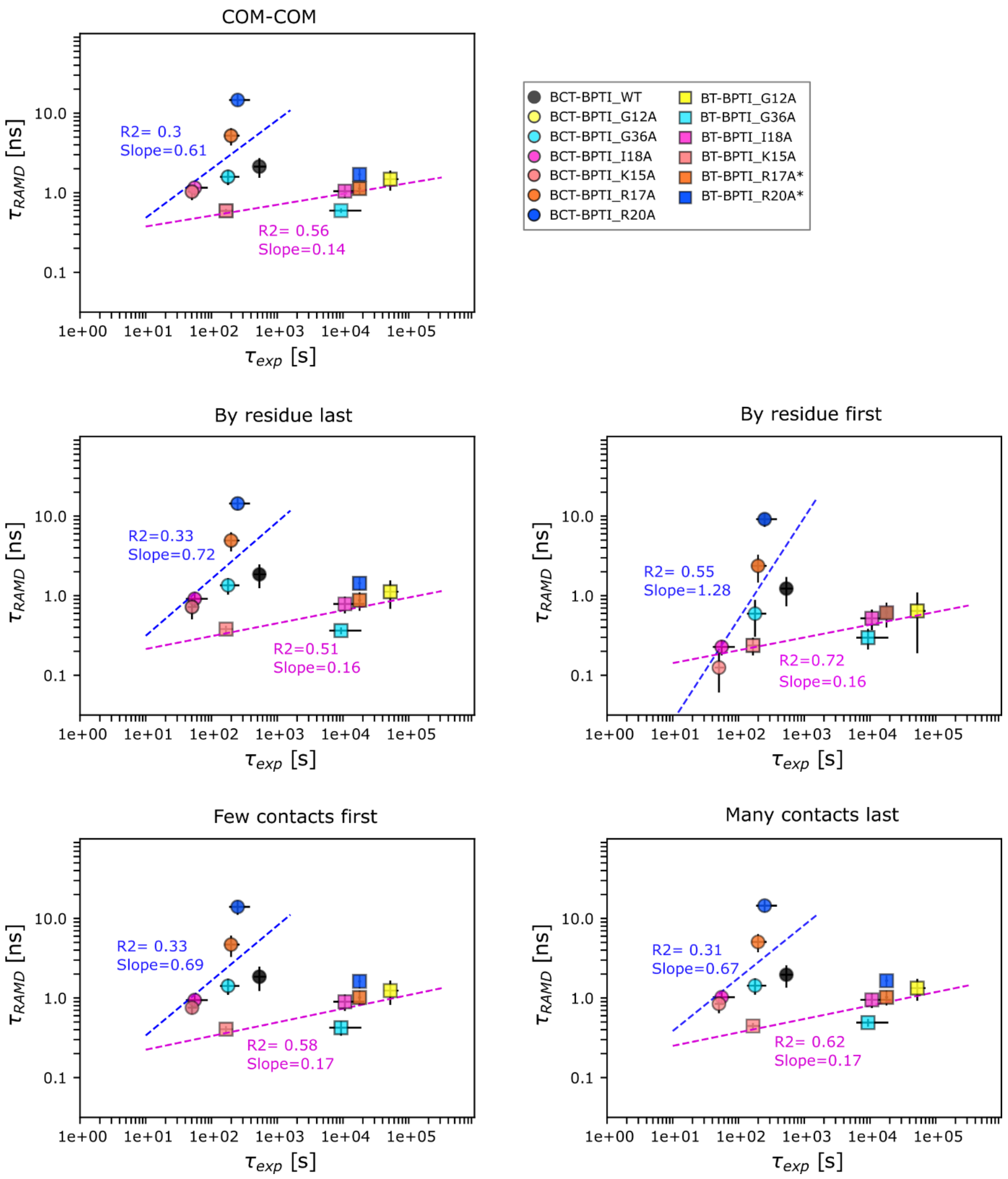


**Figure S24. RAMD residence time for the BT-BPTI and CBT-BPTI complexes computed with a random force magnitude of 19 kcal/mol/Å vs experimental residence times (inverse of $k_{off}$) for the different definitions of residence time (see Methods).** The error bars indicate standard deviations and the straight line fits are shown in magenta and blue for the BT-BPTI and BCT-BPTI complexes. BT mutants indicated with * have experimental $k_{off}$ values < $5.6x10^{-5}$ $s^{-1}$ and were not used for determining the straight line fit. The residue-first definition of the residence time gives a higher correlation and slope than the other definitions for both sets of complexes.

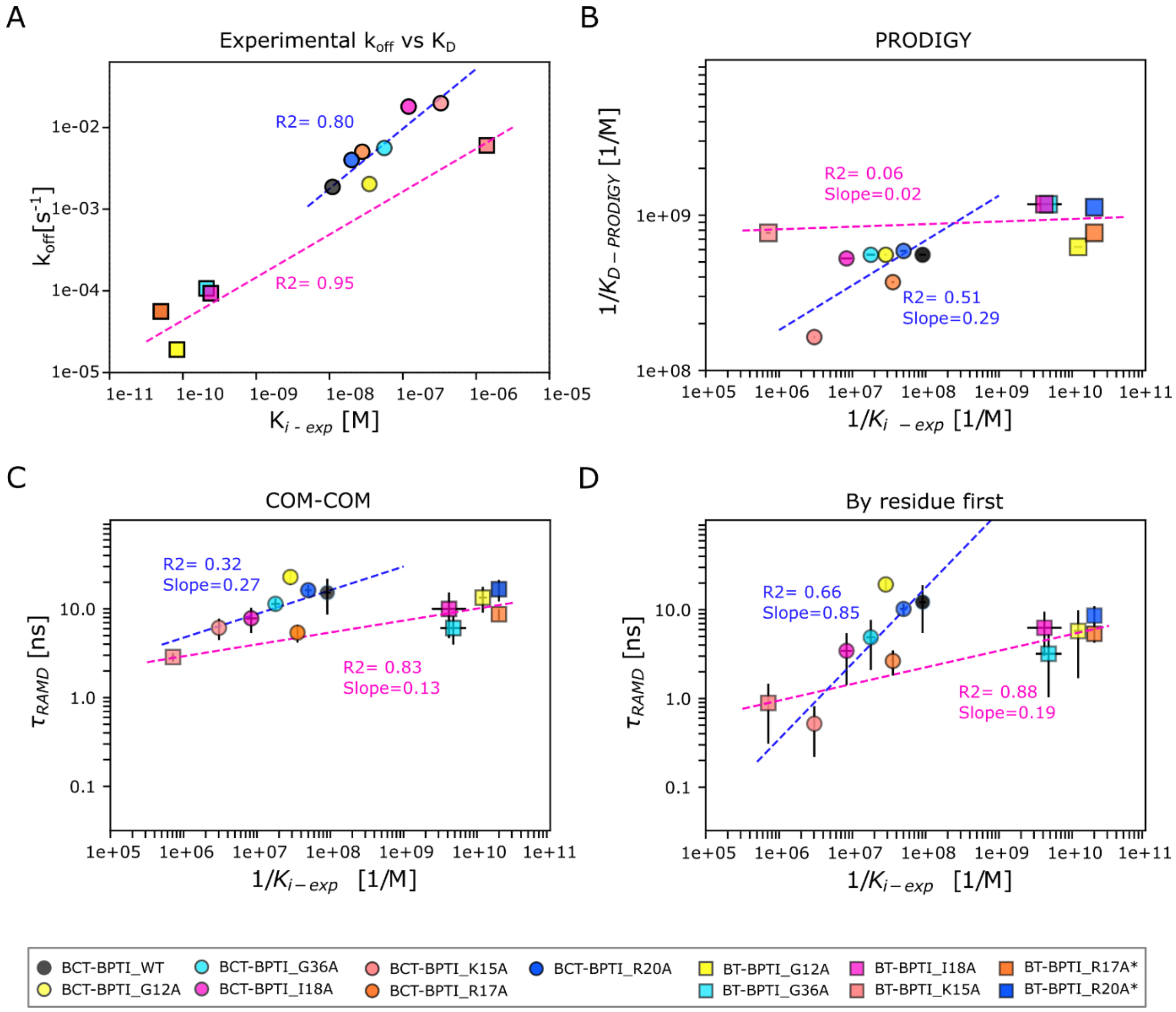


**Figure S25. Comparison with the experimentally measured inverse value of $K_i$ for the BT-BPTI and CBT-BPTI complexes**. **A)** Correlation between experimental $k_{off}$ and $K_i$ values. **B)** PRODIGY prediction of $K_i$ vs experimental values. **C-D)** RAMD residence time computed with a random force magnitude of 17 kcal/mol/Å vs experimental inverse values of $K_i$. Correlations are shown for the ''COM-COM'' and ''By residue first'' definitions of residence time (see Methods). The error bars indicate standard deviations and the straight line fits are shown in magenta and blue for the BT-BPTI and BCT-BPTI complexes. BT mutants indicated with * have experimental $k_{off}$ and $K_i$ values < $5.6x10^{-5}$ $s^{-1}$ and < $5x10^{-11}$ M, respectively, and were not used for determining the straight line fit.
